# Neuronal-Activity Dependent Mechanisms of Small Cell Lung Cancer Progression

**DOI:** 10.1101/2023.01.19.524430

**Authors:** Solomiia Savchuk, Kaylee Gentry, Wengang Wang, Elana Carleton, Belgin Yalçın, Yin Liu, Elisa C. Pavarino, Jenna LaBelle, Angus M. Toland, Pamelyn J. Woo, Fangfei Qu, Mariella G. Filbin, Mark A. Krasnow, Bernardo L. Sabatini, Julien Sage, Michelle Monje, Humsa S. Venkatesh

## Abstract

Neural activity is increasingly recognized as a critical regulator of cancer growth. In the brain, neuronal activity robustly influences glioma growth both through paracrine mechanisms and through electrochemical integration of malignant cells into neural circuitry via neuron-to-glioma synapses, while perisynaptic neurotransmitter signaling drives breast cancer brain metastasis growth. Outside of the CNS, innervation of tumors such as prostate, breast, pancreatic and gastrointestinal cancers by peripheral nerves similarly regulates cancer progression. However, the extent to which the nervous system regulates lung cancer progression, either in the lung or when metastatic to brain, is largely unexplored. Small cell lung cancer (SCLC) is a lethal high-grade neuroendocrine tumor that exhibits a strong propensity to metastasize to the brain. Here we demonstrate that, similar to glioma, metastatic SCLC cells in the brain co-opt neuronal activity-regulated mechanisms to stimulate growth and progression. Optogenetic stimulation of cortical neuronal activity drives proliferation and invasion of SCLC brain metastases. In the brain, SCLC cells exhibit electrical currents and consequent calcium transients in response to neuronal activity, and direct SCLC cell membrane depolarization is sufficient to promote the growth of SCLC tumors. In the lung, vagus nerve transection markedly inhibits primary lung tumor formation, progression and metastasis, highlighting a critical role for innervation in overall SCLC initiation and progression. Taken together, these studies illustrate that neuronal activity plays a crucial role in dictating SCLC pathogenesis in both primary and metastatic sites.

## Introduction

The nervous system is emerging as a critical component of the tumor microenvironment that regulates cancer pathobiology. Primary brain cancers such as gliomas exhibit a profound dependency on these neuronal mechanisms, both through activity-dependent paracrine signaling pathways^1–3^ as well as through direct functional integration of malignant glioma cells into electrically active neural circuits via bona fide neuron-to-glioma synapses^4–6^. While less is known about the role of neurons in brain metastases, breast cancer cells metastatic to brain were recently discovered to occupy the perisynaptic space to usurp glutamate for their growth^7^. Increasing evidence also implicates the nervous system in the regulation of many non-CNS cancers, including prostate, gastric, colon, pancreatic, breast, and skin cancers^8–17^. These studies support the emerging principle that the nervous system can profoundly influence cancer pathobiology and highlight the great number of cancers that remain to be examined from this perspective.

Small cell lung cancer (SCLC) is a lethal high-grade neuroendocrine tumor that accounts for ∼15% of all lung cancers, causes over 200,000 deaths worldwide annually^18,19^, and has a 60% chance of metastasis by the time of diagnosis, with a particular propensity to metastasize to the brain^20,21^. Experimental mouse models indicate that SCLC can originate from pulmonary neuroendocrine cells (PNECs)^22,23^, a lung epithelial cell type that resides in close proximity to nerve fibers and expresses neurotransmitter receptors^24–26^. SCLC cells have also been shown to exhibit gene expression programs that resemble those in neurons^27–29^. Higher levels of these neuronal markers correlate with shorter survival and more metastatic disease^30–32^. Recently, studies have found that these neuronal gene expression programs in SCLC are implicated in driving metastatic progression by facilitating interactions with astrocytes in the brain microenvironment^33^. Yet, whether neuronal activity may be a regulator of progression and metastatic ability is not yet understood. We thus hypothesized that nervous system-SCLC interactions may drive tumor progression. Here, we investigate the role of neurons and neuronal activity on the growth, progression and spread of primary SCLC in the lung and SCLC in the brain.

### Neuronal activity drives the progression of SCLC brain metastases

To evaluate interactions of neurons with SCLC cells metastatic to the brain, we first analyzed metastatic SCLC brain tissue samples from five patients. Regions of the tumor mass demonstrated extensive axons intermingled with malignant cells as determined by immunohistochemical neurofilament staining (Fig. 1a, Extended Data Fig. 1a), while other regions exhibited little to no axonal infiltration. Quantifying the proliferation index of SCLC cells in regions of the mass infiltrated by axons compared to regions with a dearth of axons, we found that malignant cells that were closer to axons (<100 μm) exhibited increased rates of proliferation (Fig. 1b), suggesting a possible functional role of neuron-SCLC interactions.

**Figure 1.**
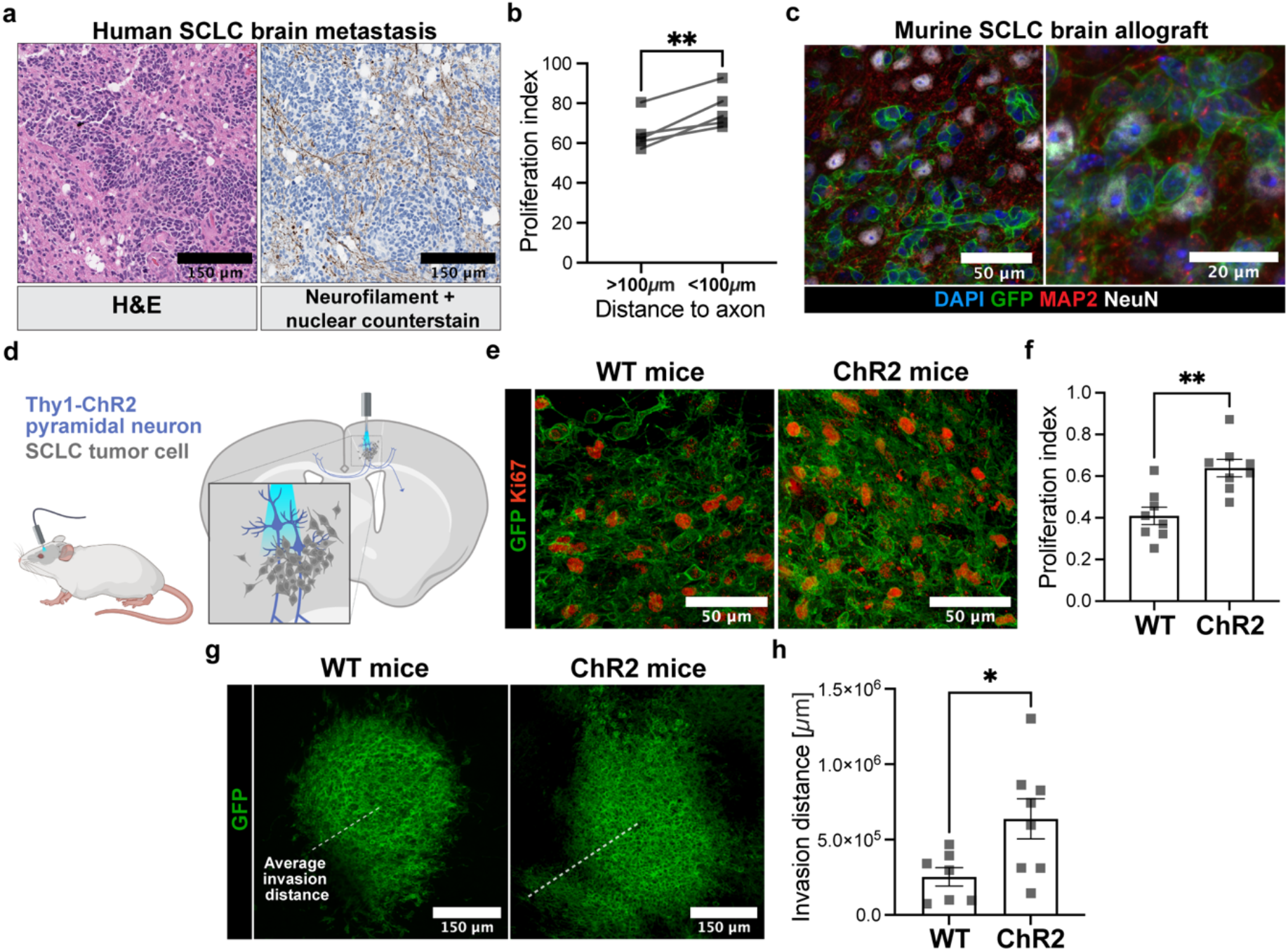
Neuronal activity promotes metastatic SCLC progression in the brain. **a**, Representative histology of tissue taken from human SCLC brain metastases. Left, H&E staining; right, immunohistochemistry for neurofilament (brown) with nuclear counterstain (blue). Scale bar = 150 μm. **b**, Proliferation index in regions of human SCLC brain metastases quantified either less than or greater than 100 μm from axons immunostained for neurofilament (n=5 patients). **c**, Representative immunofluorescent images of murine SCLC brain allografts (GFP, green) with neurons labeled with MAP2 (red) and NeuN (grey). Scale bar = 50 μm (left), 20 μm (right). **d**, Experimental paradigm for *in vivo* optogenetic stimulation of Thy1::Chr2 pyramidal premotor cortical projection neurons in awake behaving mouse with SCLC tumor allografted into the M2 cortex. **e**, Representative immunofluorescent images of murine SCLC brain tumors (GFP, green) allografted in cortical region of wild-type (WT) or Thy1::ChR2 (ChR2) animals. Proliferating cells labeled with Ki67 (red). Scale bar = 50 μm. **f**, Quantification of data in **e**, illustrating increased proliferation of tumors in optogenetically stimulated animals (n=8 WT, n=8 ChR2 animals). **g**, Immunofluorescent images demonstrating invasion area of SCLC brain allografts of WT or ChR2 animals. Scale bar = 150 μm. **h**, Quantification of data in **g**, illustrating increased invasion distance after optogenetic stimulation (n=7 WT, n=8 ChR2 animals). Data are mean ± s.e.m. **P< 0.01, *P< 0.05, two-tailed paired t-test for **b**, two-tailed unpaired t-test for **f, h**.

As there is no reliable model of spontaneous brain metastasis in any of the existing genetic mouse models of SCLC^34^, we used murine intracranial allografts to test potential functional influences of neuronal activity on progression of SCLC metastases in the brain. This permitted us to place the malignant cells into a specific region of interest, allowing direct manipulation of distinct neural circuits involving the tumor. It is important to note the caveat that this is not a model of metastatic initiation, but rather one of growth of an established metastatic tumor. These allografts exhibited presence of neurons in the tumor mass, similar to the human samples described above, especially in areas of the tumor periphery (Fig. 1c). To examine the role of neuronal activity on metastatic SCLC tumor progression, we employed *in vivo* optogenetic techniques in freely behaving mice. Here, murine SCLC cells (16T) were allografted intracranially into the premotor cortex of mice expressing the blue-light-sensitive opsin channelrhodhopsin (ChR2) in Thy1+ deep layer cortical projection neurons or wild-type (WT) littermate controls (Fig. 1d). After a period of engraftment, optogenetic ferrules were placed over the premotor cortex and mice were stimulated with blue light (473 nm, 20 Hz; cycles of 30 s on/90 s off over 30 min). Successful stimulation of cortical neurons on Thy1::ChR2 animals was verified by the observance of a complex motor behavioral output, unilateral ambulation. Light stimulation had no behavioral effect in identically manipulated allografted WT mice. Assessed histologically at 24 hours after optogenetic stimulation of cortical neuronal activity, SCLC cells exhibited a robust increase in proliferation rate (∼40% Ki67+ SCLC cells in mock-stimulated, WT mice, ∼60% Ki67+ SCLC cells in optogenetically stimulated ChR2-expressing mice; Fig. 1e-f). Further, there was a clear increase in the spread of SCLC cells in the brain, with more cells migrating outside of the core of the tumor in mice with optogenetically elevated neuronal activity (Fig. 1g-h). Taken together, these results indicate that cortical neuronal activity can promote the proliferation and invasion of SCLC cells in the brain.

### Neuron-SCLC interactions

To further probe the interactions between SCLC and neuronal activity, we utilized co-culture systems in which either cortically derived murine neurons or human iPSC-derived neurons were cultured together with either murine SCLC (16T^28^, subtype SCLC-A) or human SCLC (H446^35^, subtype SCLC-N) cells. In all cases, the presence of active neurons significantly increased the rate of proliferation of these clinically and molecularly distinct subtypes of SCLC cells in co-culture (Fig. 2a-c, Extended Data Fig. 2a-c). This effect was abrogated with the addition of tetrodotoxin (TTX), a voltage-gated sodium channel blocker that inhibits action potentials (Fig. 2b-c, Extended Data Fig. 2b-c). To determine whether key activity-dependent paracrine factors that we previously have shown to robustly affect glioma growth^1,3,5,36^ similarly induce proliferation in SCLC, we added recombinant NLGN3 (100nM) and BDNF (100nM) to murine SCLC cells *in vitro*. Neither of these paracrine factors were found to affect the proliferation of the lung cancer cells, in contrast to a clear growth-promoting effect in patient-derived glioma cultures (Extended Data Fig. 2d-g), suggesting that the observed proliferative effects of neuronal activity are driven by distinct mechanisms in the two tumor types.

**Figure 2.**
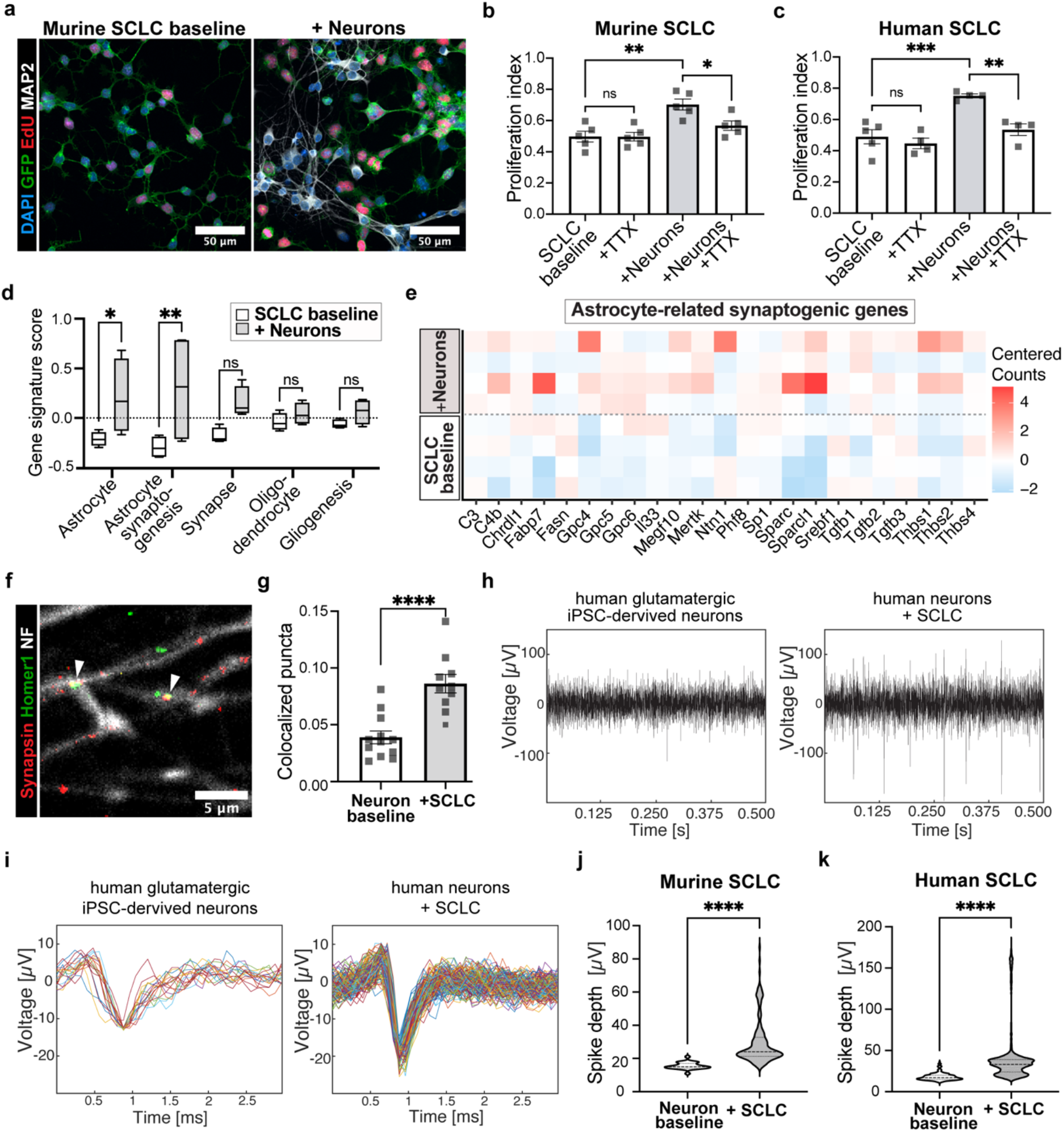
Reciprocal effects of neuron-SCLC interactions. **a**, Representative immunofluorescent images of murine SCLC cells (16T; green) co-cultured with primary cortical neurons labeled with MAP2 (white). Proliferative cells are labeled with EdU (red). Scale bar = 50um. **b**, Quantification of proliferative index of murine SCLC cells co-cultured with primary cortical neurons with or without addition of 1 μM TTX (n=5 coverslips per condition). **c**, as in **b**, for human SCLC cells (H446) co-cultured with primary cortical neurons (n=4 coverslips per condition). **d**, Gene signature scores generated from centered RNA-seq gene expression data from cultured murine and human SCLC cells at baseline or isolated from neuronal co-cultures (n=4 independent samples per condition; for gene sets used to generate the signature see Extended Data Table 1). **e**, Heatmap demonstrating centered counts of astrocyte-related synaptogenesis genes from in-vitro cultured murine and human SCLC cells at baseline or isolated from neuronal co-cultures (n=4 independent samples per condition; see Extended Data Table 1). **f**, Representative confocal image of human iPSC-derived neurons co-cultured with murine SCLC cells (16T). Arrowheads indicate colocalized pre-synaptic puncta (Synapsin, red) and post-synaptic puncta (Homer 1, green) detected along neuronal processes. Neurofilament (white) is used to visualize neuronal fibers. Scale bar = 5 μm. **g**, Quantification of the number of colocalized pre- and post-synaptic puncta on neuronal processes at baseline vs. co-cultured with SCLC cells defined per 10 μm of neurofilament length (n=12 coverslips in neuron baseline condition, n=10 coverslips in SCLC co-culture condition). **h**, Representative recording over 500 ms demonstrating spike number in human iPSC-derived neurons at baseline (left) vs. co-cultured with murine SCLC cells (16T; right) using multielectrode array recording systems (n=6 per condition). **i**, Representative trace illustrating increased spike amplitude in neurons co-cultured with SCLC cells compared with baseline. **j**, Quantification of spike number and amplitudes from **h and i** shown as violin plot (n=24 spikes in neuron baseline and n=168 spikes in SCLC co-cultured neurons). **k**, As in **j**, but for human SCLC cells (H446) (n=363 spikes in neurons baseline and n=2721 spikes in SCLC co-cultured neurons). Data are mean ± s.e.m. for **b, c, g**, box-and-whiskers plot for **d**, violin plot for **j, k**; analysis with 2-way ANOVA for **b, c, d**, two-tailed unpaired t-test for **g**, Mann-Whitney test for **j, k**; ****P<0.0001, ***P<0.001, **P< 0.01, *P< 0.05

We next examined the transcriptional phenotypic changes in SCLC cells following exposure to neurons by isolating either the GFP-labeled murine or human SCLC cells that were co-cultured with neurons using flow cytometry. These malignant cells were then subjected to RNA-seq analysis. To ensure the lack of neuronal contamination, housekeeping genes that are typically highly expressed in neurons were specifically queried. We found no change in the expression of these genes in SCLC cells isolated from either pure SCLC cultures, or those isolated from neuronal co-cultures (Extended Data Fig. 3a). In line with the proliferative effect described above, we found that upregulation of a cell proliferation signature was a prominent effect of neuronal co-culture across both human and murine SCLC cells (Extended Data Fig. 3b). When comparing murine SCLC cells transplanted to the flank or to the brain of mice^33^, we found a substantial number of SCLC genes upregulated in the context of the brain that overlapped with the genes found to be upregulated in the presence of neuronal co-cultures (Extended Data Fig. 3c). Gene ontology analyses of these SCLC genes with enriched expression in the presence of neurons similarly revealed a dominant presence of a proliferative signature, illustrating again that the neural microenvironment promotes SCLC proliferation (Extended Data Fig. 3d).

In addition to the upregulation of proliferative signatures, neuronal co-culture increased the expression of a gene signature associated with an astrocytic phenotype^37–39^ in SCLC cells (Fig. 2d, Extended Data Fig. 3e, Extended Data Table 1), reminiscent of the astroglial-like SCLC population that has been identified in previous studies^40^. Upregulation of this gene set was also accompanied by a shift in SCLC cell morphology from exhibiting long thin processes to shorter processes with fewer ‘spine-like’ structures (Extended Data Fig. 4a-e). This shift was specific to SCLC cells, as patient-derived glioma cells (SU-DIPG-XIIIFL and SU-DIPG-VI) demonstrated the opposite effect, exhibiting longer thin processes in the presence of neurons (Extended Data Fig. 4f-j). SCLC cells in isolation typically exhibit a neuron-like morphology^28,29,41^, therefore this shift towards an astrocyte-like phenotype suggested another functional influence of neurons on SCLC.

Further analyzing the changes in gene expression, we found that in both human and murine SCLC, exposure to neurons increased the expression of astrocyte-associated synaptogenic genes (Fig. 2d, e). Astrocytes are known to play a central role in the form and function of neuronal circuits by secreting synaptogenic factors, amongst other functions^42–47^. As cancers metastatic to brain are often associated with seizures^48–51^ and astrocyte-like glioma cells have been known to create a hyperexcitable neuronal microenvironment through the secretion of glutamate and synaptogenic factors^52–54^, we next assessed whether these astrocyte-like SCLC cells in the brain could similarly - and reciprocally - affect neurons in the microenvironment. To test the contribution of SCLC cells to neuronal hyperexcitability, we first quantified co-localization of presynaptic puncta (synapsin) with postsynaptic puncta (Homer1) on neuronal filaments in our SCLC-neuron co-cultures. We found a marked increase in the number of synapses on neurons co-cultured with SCLC cells compared to their baseline (Fig. 2f, g). Additionally, using multielectrode array systems, we found that in the presence of SCLC cells, these co-cultured neurons exhibited elevated activity measured both by increased spike number and amplitude (Fig. 2h-k). This further implicates SCLC cells in modulating neuronal activity—suggesting that malignant SCLC cells are capable of remodeling the tumor microenvironment in the brain. To determine whether this effect was due to any cell-intrinsic changes in pyramidal neurons associated with SCLC, we performed *in slice* whole cell patch clamp recordings of pyramidal neurons either integrated in regions of the tumor or in the contralateral (control, non-tumor infiltrated) hippocampus. Between the two regions, no distinct differences in cellular properties were noted (Extended Data Fig. 5). These observations are consistent with the idea that malignant astrocyte-like SCLC cells may contribute to driving hyperexcitability in the tumor microenvironment through increased synaptogenesis^53,55^, thereby reinforcing neuron-SCLC interactions.

Carcinoma cells metastatic to brain in breast and non-small cell lung cancer models establish gap junction-mediated connections to astrocytes to enable astrocyte-cancer cell signaling that promotes brain metastasis growth^56^. Similarly, astrocyte-like glioma cells form a gap-junction coupled network between astrocyte-like glioma cells and between glioma cells and true astrocytes for the purposes of intercellular communication and therapeutic resistance^57,58^. Here we find that neurons influence SCLC cells to assume a more astrocyte-like phenotype, which could enable similar, growth-promoting gap-junction-mediated signaling. To assess the functional role of SCLC gap junctions to metastatic outgrowth in the brain, tumor-bearing mice were treated with meclofenamate, a clinically available gap-junction blocker. Meclofenamate-treated animals demonstrated reduced growth of SCLC in the brain, as measured by *in vivo* bioluminescent imaging of tumor burden, suggesting a growth-promoting role for such gap junctional connections (Extended Data Fig. 6a-b).

### Neuronal activity-dependent currents in SCLC cells

In addition to upregulation of the astrocyte-associated genes, SCLC cells co-cultured with active neurons exhibit increased expression of a multitude of neurotransmitter receptor and synapse-related genes (Fig. 3a, Extended Data Fig. 3f, Extended Data Fig. 7a-c, Extended Data Table 1). To examine whether these neurotransmitter receptors are functionally relevant to the increased proliferation observed in response to neuronal activity, we added specific neurotransmitter receptor inhibitors, MK801 (NMDA receptor inhibitor) and picrotoxin (GABA receptor inhibitor), to neuron-SCLC co-cultures and quantified SCLC proliferation. The addition of each of these inhibitors diminished the neuronal activity-induced proliferation of the SCLC cells (Fig. 3b-c), indicating involvement of glutamatergic and GABA-ergic signaling in the SCLC proliferative response to neuronal activity.

**Figure 3.**
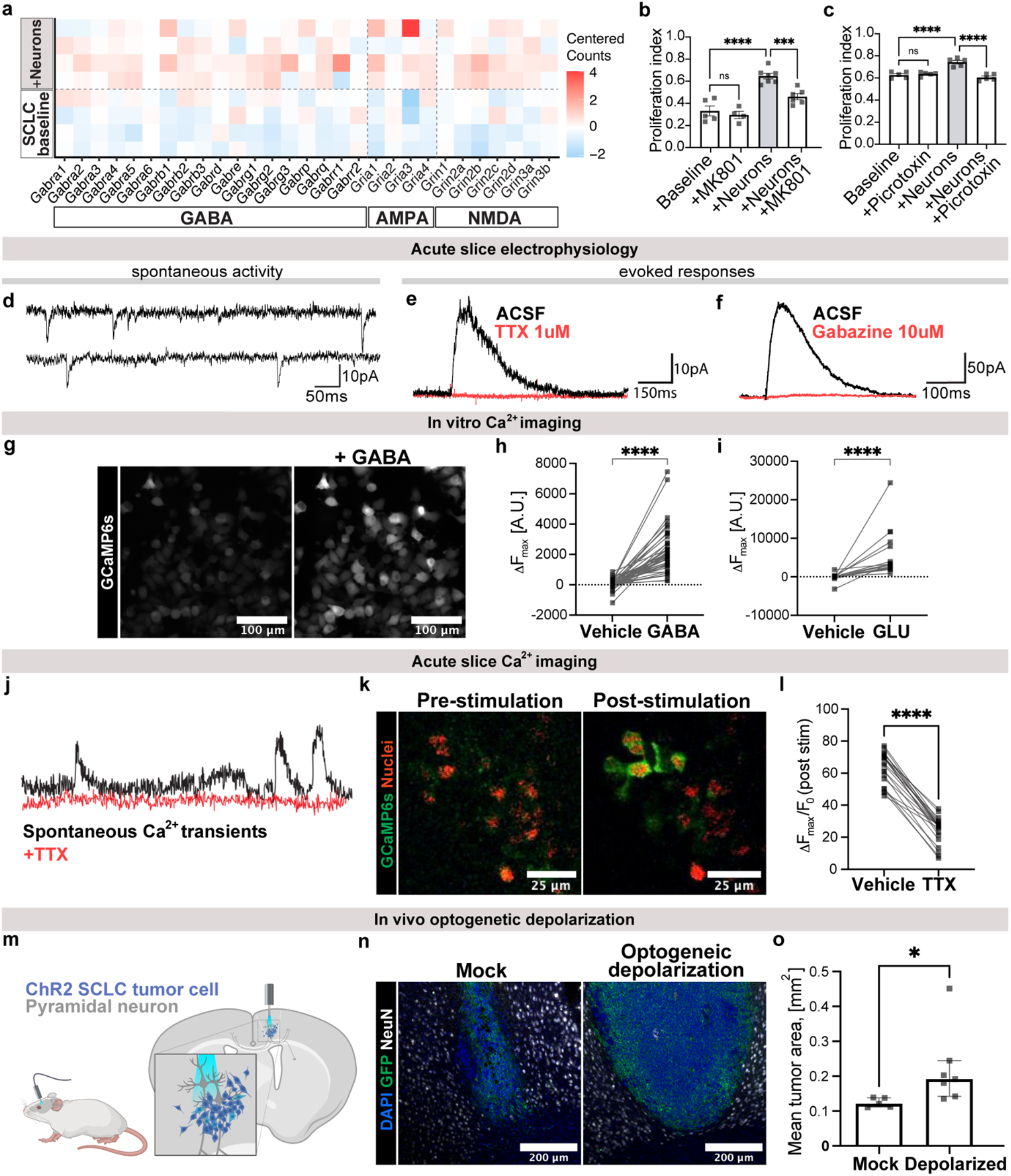
Neuronal activity-mediated currents in drive progression in SCLC. **a**, Heatmap demonstrating centered counts of selected ion channel genes from murine and human SCLC cells at baseline or isolated from neuronal co-cultures (n=4 independent samples in each condition, for gene list see Extended Data Table 1). **b**, Quantification of proliferative index of murine SCLC cells co-cultured with neurons with or without addition of 75 μM picrotoxin (n=5 coverslips per condition). **c**, Quantification of proliferative index of murine SCLC cells co-cultured with neurons with or without addition of 50 μM MK801 (n=5 coverslips per condition). **d**, Representative recordings of spontaneous currents in allografted SCLC cells (n =19/86 cells). **e**, Representative traces of neuronal-activity dependent evoked SCLC currents before (black) and after (red) application of 0.5 μM TTX (n=15/22 cells). **f**, As in **e**, but with 10 μM gabazine. **g**, Representative calcium imaging of GCaMP6s-containing SCLC cells at baseline (left) and after application of 1 mM GABA. Scale bar = 100 um. **h**, Quantification of GCaMP6s fluorescence in individual SCLC cells (from **l**) in response to administration of 1 mM GABA (n=44 cells). **i**, As in **h**, but with administration of 1 mM glutamate (n=19 cells). **j**, Two-photon in slice calcium imaging of GCaMP6s-expressing SCLC cells in hippocampal allografts. Representative trace of spontaneous activity as measured by changes to GCaMP6s fluorescence in SCLC cells with (red) or without (black) administration of 0.5 μM TTX. **k**, Two-photon in situ calcium imaging in GCaMP6s-expressing SCLC cells in hippocampal allografts with Schaffer collateral stimulation (*n* = 6 slices, 4 mice). Representative frames shown before (left) and after (right) stimulation. Red denotes SCLC cancer cell tdTomato nuclear tag; green denotes SCLC GCaMP6s. Scale bar = 25 μm. **l**, Quantification of GCaMP6s fluorescence in individual SCLC cells in response to electrical stimulation of CA1 Shaffer collateral axons with or without administration of 0.5 μM TTX (n=22 cells). **m**, Experimental paradigm for in-vivo optogenetic depolarization of intracranial allografts of channelrhodopsin-2 (ChR2)-expressing SCLC cells. **n**, Representative confocal images of ChR2-expressing SCLC allografts after mock or blue-light-induced depolarization. Neuronal nuclei are labeled with NeuN (white), tumor cells in GFP (green). Scale bar = 200 um. **o**, Quantification of mean tumor area from **n** (n=5 mock, n=7 depolarized mice). Data are mean ± s.e.m. for **b, c**, median ± IQR for **o**; analysis with two-tailed unpaired t-test for **b, c**, paired t-test for **h, i, l**, Mann Whitney test for **o;** ****P< 0.0001, ***P< 0.001, **P< 0.01, *P< 0.05.

Having established the functional role of specific neurotransmitter receptors in neuron-SCLC growth-promoting interactions, we next assessed electrophysiological responses of SCLC to neuronal activity. GFP-labeled SCLC 16T cells were allografted into the well-mapped CA1 region of the hippocampal circuit (Extended Data Fig. 7d-e). After a period of engraftment and growth, acute hippocampal slices were prepared for whole-cell patch-clamp recordings of GFP+ SCLC cells. Electrophysiological recordings in voltage clamp at −70 mV of individual SCLC cells revealed the presence of spontaneous currents, resembling excitatory glutamatergic currents, in a subpopulation of cells (∼22% of SCLC cells; Fig. 3d).

To assess for neuronal activity-dependent evoked responses, we stimulated CA3 Schaffer collateral and commissural axons, the afferent input to the CA1 region, while simultaneously recording from SCLC cells in CA1. At voltage clamp of −70 mV to minimize GABA currents, evoked responses were only very rarely seen. When held at 0 mV to minimize glutamatergic currents, a high proportion of SCLC cells (∼68%) exhibited GABA-ergic currents in response to neuronal activity. These currents were blocked with the addition of TTX (Fig. 3e) or gabazine (Fig. 3f), a GABA-receptor inhibitor. These observations of SCLC GABAergic currents are consistent with the findings that pulmonary neuroendocrine cells secrete GABA and express GABA receptors in the normal lung^59,60^. Together, these electrophysiological studies reveal activity-induced and neurotransmitter-mediated signaling from neurons to SCLC cells in the brain.

We next visualized these currents in SCLC cells using calcium imaging. Applying each neurotransmitter individually to SCLC cells engineered to express the genetically encoded calcium indicator GCaMP6s *in vitro*, we found that both GABA and glutamate exposure elicits calcium transients (increased GCaMP6s fluorescence) in these malignant cells (Fig. 3g-i). We then performed *in slice* two-photon calcium imaging of GCaMP6s-expressing SCLC cells in the hippocampus. Similar to what was observed using electrophysiology, both spontaneous and axonal stimulation-evoked, neuronal activity-dependent calcium transients in the SCLC cells were observed (Fig. 3j-k). These transients were blocked by the addition of tetrodotoxin, indicating their reliance upon neuronal activity (Fig. 3l).

### SCLC membrane depolarization regulates growth

Given the activity-dependent SCLC currents and consequent calcium transients described above, we next sought to understand whether membrane depolarization provides a functional benefit to SCLC tumors in the brain. We employed optogenetics to directly depolarize SCLC cells engineered to express channelrhodopsin-2 (ChR2). These cells were allografted into the mouse cortex (Fig. 3m). After a period of engraftment, blue-light was delivered via a fiber optic placed at the surface of the brain to directly depolarize these ChR2-expressing SCLC tumor cells *in vivo* (473 nm, 10 Hz, 30 s on / 90 s off for 30 min). Control animals were identically manipulated with mock (no blue light) stimulation. Functionality of the ChR2 construct in SCLC cells was confirmed with patch clamp electrophysiology (Extended Data Fig. 7f-g). Membrane depolarization resulted in a robust growth effect in SCLC, with the overall size of the tumor approximately doubling after three stimulation sessions compared to identically manipulated, mock-stimulated control mice (Fig. 3n-o). Taken together, these studies illustrate that SCLC cells in the brain utilize neuronal activity and the resulting SCLC membrane depolarization to fuel growth and progression of the metastatic tumor.

### The role of innervation in the primary lung tumors

Given the substantial role of neurons in the progression of SCLC brain metastases described above, we next examined the role of innervation in the context of primary lung SCLC initiation and metastasis. Analysis of human primary SCLC samples taken from patient lung biopsies^61^ revealed the expression of several neurotransmitter receptor genes, consistent with the idea that primary-site SCLC cells possess the ability to respond to neuronal cues (Fig. 4a, Extended Data Fig. 8a, Extended Data Table 1). To further investigate the influence of neural input on SCLC pathobiology in the lung, we utilized the *Trp53*^*flox/flox*^, *Rb1*^*flox/flox*^, *p130*^*flox/flox*^ knockout (RPR2), luciferase (luc)-expressing SCLC genetic mouse model^62^. Tumors were induced in 8-week-old RPR2-luc mice by intratracheal administration of Adeno-CMV-Cre. These mice form spontaneous SCLC tumors that recapitulate the genetics, histology, therapeutic response, time course of progression, and highly metastatic nature of the human disease and allow for the isolation of primary tumors and metastases (e.g. to liver) directly from their native microenvironment^63–65^. Examination of SCLC tumors taken from the mouse lung revealed innervation of malignant tissue by various nerve types, including parasympathetic (labeled by vesicular acetylcholine transporter protein), sympathetic (labeled by tyrosine hydroxylase protein), and sensory (labeled by myelin basic protein) nerve fibers (Fig. 4b). These findings are consistent with the known innervation patterns of normal lung largely from vagal nerve parasympathetic efferents and sensory afferents, together with sympathetic and sensory fibers from the thoracic spinal cord sympathetic chain and dorsal root ganglions, respectively^66^. An abundance of nerve fibers were also evident in the vicinity of tumors metastatic to the liver in these animals (Extended Data Fig. 8b).

**Figure 4.**
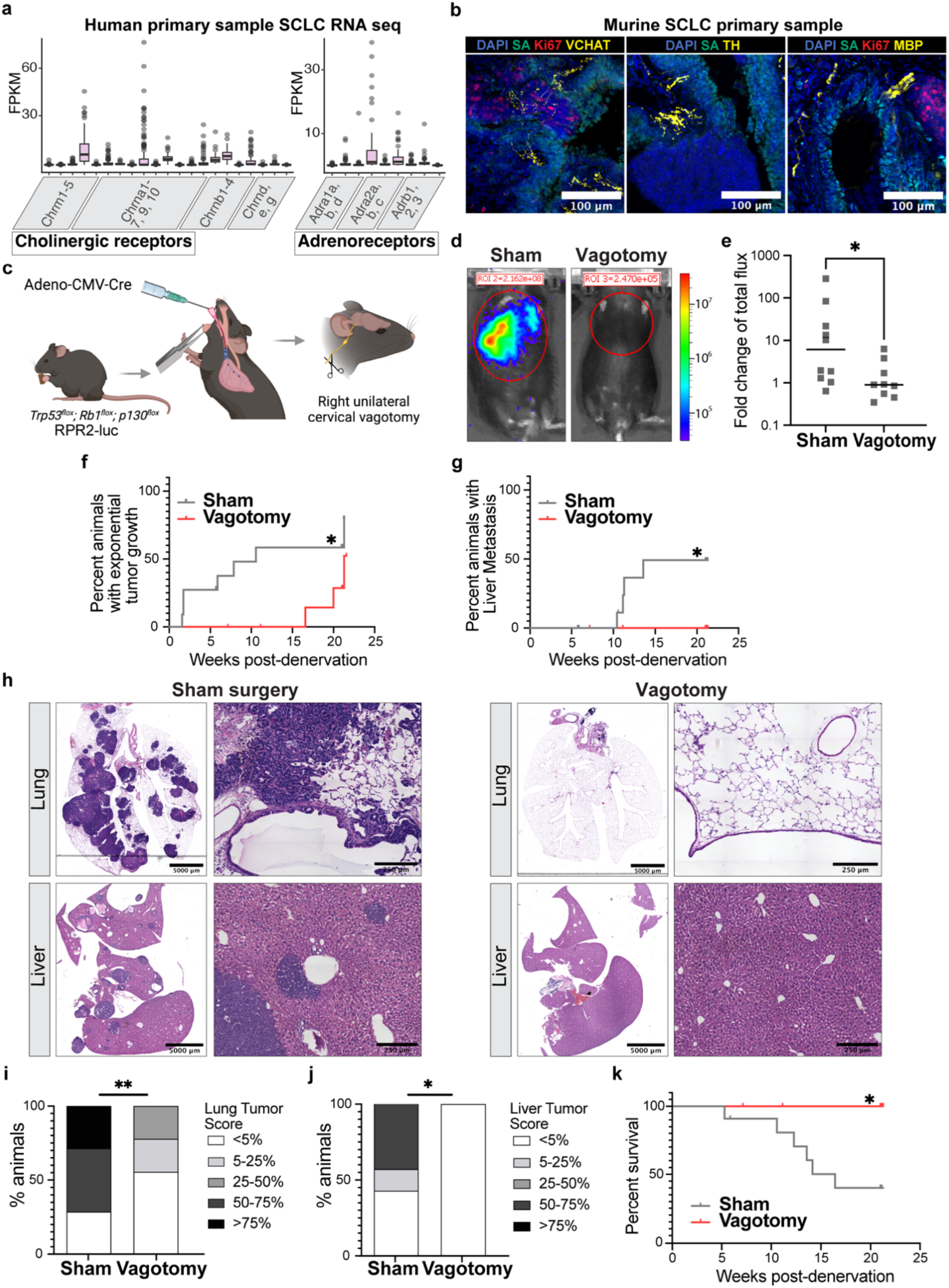
Vagal nerve innervation is critical for primary SCLC tumor formation and metastasis. **a**, Quantification of key neurotransmitter receptor gene expression in human samples of primary SCLC (n=81, for gene list see Extended Data Table 1). **b**, Visualization of tumor innervation in primary lung tumor samples taken from genetic model of spontaneously forming SCLC. Streptavidin (SA, green) used to mark airways^67^; proliferative cells are labeled with Ki67 (red) to help identify tumors. Vesicular acetylcholine transporter (VAChT), tyrosine hydroxylase (TH), and myelin basic protein (MBP) (yellow) are used to visualize parasympathetic, sympathetic, and myelinated sensory populations of nerve fibers in left, middle and right image, respectively. Scale bar = 100 μm. **c**, Experimental paradigm for unilateral cervical vagotomy in genetic model of spontaneously forming SCLC (RPR2-luc). **d**, Representative IVIS image of RPR2-luc animals 10 weeks after sham (left) or vagotomy (right) procedure. Heat map, photon emission. **e**, IVIS bioluminescence analysis of overall tumor growth in SCLC primary tumors measured at 10 weeks after vagotomy procedure (n=10 sham and n=9 vagotomy animals from 2 independent cohorts). Data represented as fold change in total flux. **f**, Time course of tumor growth in RPR2-luc mice as measured by IVIS bioluminescence imaging after sham-operation (grey) or vagotomy procedure (red); (n=11 sham and n=9 vagotomy animals). **g**, Time course of liver metastasis onset in RPR2-luc mice as detected by IVIS bioluminescence imaging after sham-operation (grey) or vagotomy procedure (red; n=11 sham and n=9 vagotomy animals). **h**, Representative H&E of lungs (top) and livers (bottom) isolated from sham-operated and denervated (vagotomy) RPR2-luc mice. Scale bar = 5000 μm (left image) and 250 μm (right image). **i**, Quantification of lung tumor score (percentage of the organ occupied by the tumor) in sham-operated and denervated (vagotomy) RPR2-luc mice (n = 7 sham, n=9 vagotomy animals). **j**, as in **i**, for quantification of liver tumor score (n = 7 sham, n=9 vagotomy animals). **k**, Kaplan–Meier survival curve of RPR2-luc mice after either sham-operation (grey) or denervation (vagotomy; red); (n=11 sham and n=9 vagotomy animals). Data are box-and-whisker plots for **a**, median for **e**; analyzed with Mann Whitney test for **e**, Gehan-Breslow-Wilcoxon test for **f**, Log-rank (Mantel-Cox) test for **g, k**, Fisher’s exact test for **i, j**. **P< 0.01, *P< 0.05.

To modulate innervation to the lung, we performed unilateral cervical vagotomy in this genetically engineered SCLC mouse model (Fig. 4c). Specifically, cervical vagotomy entails the direct transection of vagal sensory fibers and diminished input to the post-ganglionic parasympathetic fibers via the direct transection of pre-ganglionic vagal fibers. Vagotomies or sham surgeries were performed approximately two months after the intratracheal administration of the Adeno-CMV-Cre vector but prior to tumor initiation. Animals were then followed for the next 25 weeks using *in vivo* bioluminescent imaging. Both sham-manipulated and denervated animals tolerated the procedure well, displaying no postoperative weight loss or sickness behaviors (Extended Data Fig. 8c).

By 10 weeks post-denervation, a marked difference was evident in the overall tumor burden in sham-manipulated (control) and denervated groups (Fig. 4d-e). Longitudinal *in vivo* bioluminescent imaging revealed that both the initiation of primary lung tumors and the appearance of any metastasis were significantly delayed or not detected in the vagotomy cohort, a difference that continued throughout the remainder of the experiment (Fig. 4f-g). At the endpoint (∼6 months post-vagotomy), these differences in overall tumor burden were histologically validated. While sham-manipulated animals demonstrated an abundance of tumor sites throughout all lobes of the lung, the denervated animals illustrated minimal to no tumor burden (Fig. 4h-i). As the liver is the first site of metastasis in this mouse model, livers were carefully evaluated and found to be free of tumor in all denervated mice as assessed by both histological analysis (Fig. 4h-j), as well as gross analysis of the surface of the organ (Extended Data Fig. 8d). These results were independently evaluated and validated by a board-certified pathologist. A caveat to note is that the lack of metastatic spread to the liver is influenced, at least in part, by the marked reduction in lung tumor burden in vagotomized mice. Finally, as overall tumor burden was greatly reduced by vagotomy, we observed a striking survival benefit for the mice that had been denervated, with all denervated mice surviving the full length of the experiment compared to a median survival of 16 weeks for sham manipulated mice (Fig. 4k). In contrast, when vagotomy was performed after tumor initiation in an independent cohort of animals, such reduction of tumor burden and survival benefit were not observed (Extended Data Fig. 8e-f). Taken together, these findings suggest that vagal innervation of the primary tumor site (lung) is required for SCLC initiation and progression, but not maintenance of already established tumors.

## Discussion

The influences of neuronal activity on SCLC primary lung tumor initiation, metastatic spread to the liver, and the progression of brain metastases demonstrated here adds to the growing evidence highlighting the importance of neurobiology in cancer. These findings indicate that metastatic SCLC cells that have colonized the brain take advantage of neuronal activity to enhance growth and invasion, while reciprocally increasing neuronal excitability and activity. Developing a longitudinal understanding of the roles these neural pathways play across tumor initiation, growth, and metastases will be critical to developing novel therapies that target nervous system interactions with SCLC.

In the future, further elucidation of the specific molecular signaling cascades that mediate this activity-driven progression of SCLC may reveal therapeutic vulnerabilities. The growth promoting effect of SCLC membrane depolarization – reminiscent of similar effects in glioma^4^ – warrants further investigation of the voltage-sensitive mechanisms of neuron-SCLC crosstalk. The stark effect of denervation on pulmonary SCLC pathobiology raises questions about the distinct involvement and relative contributions of the different axonal subpopulations within the vagus nerve, the answers to which would provide potential therapeutic targets such as neurotransmitter receptors and inform potential translation of these findings to the bedside. Further, evaluating the possible effects of neuronal activity on other cells in the lung tumor microenvironment, including immune, vascular, and other stromal cells may elucidate additional indirect effects of nervous system activity on SCLC pathogenesis. SCLC is a disease with limited treatment options and poor prognosis. Thus, disrupting the functional interactions between neurons and SCLC represents a promising therapeutic avenue to explore for this lethal cancer.

## Extended figures

**Extended Data Fig. 1.**
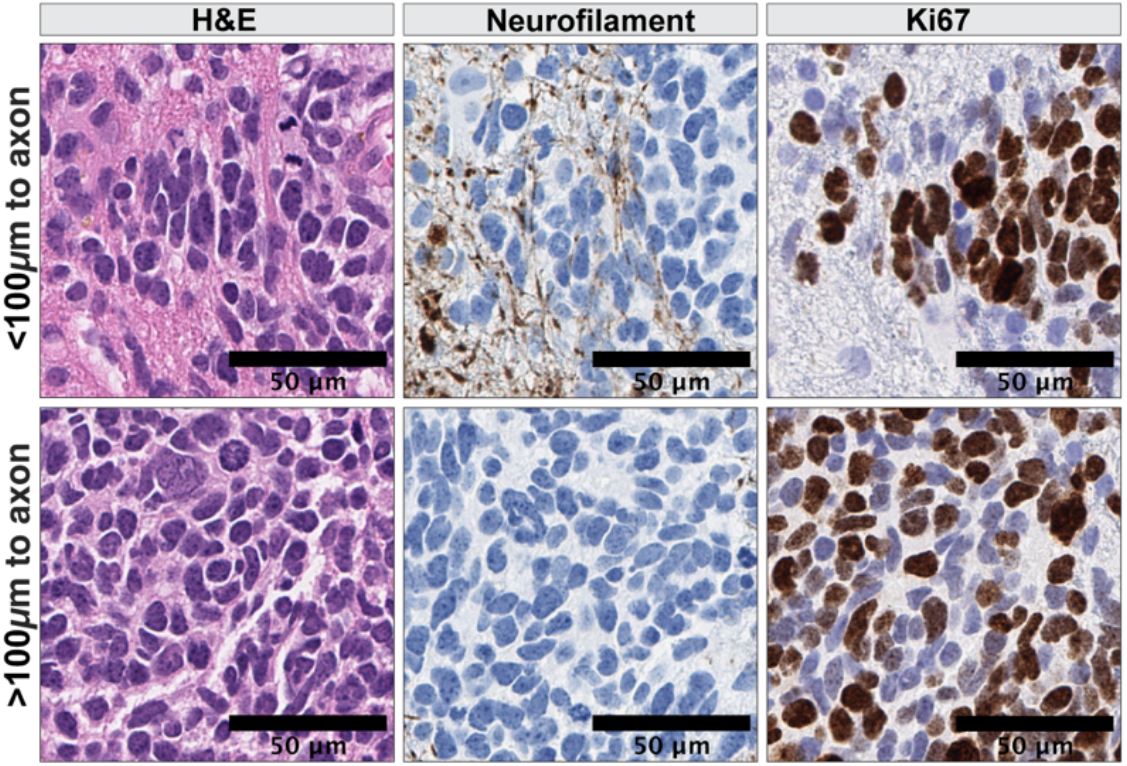
Histology of metastatic SCLC patient tissue taken from the brain. Representative H&E (left), neurofilament (middle) and ki67 (right) immunohistochemistry of human SCLC brain metastasis in regions within (top) and outside (bottom) 100 μm distance from neurofilament. Scale bar = 50 μm.

**Extended Data Fig. 2.**
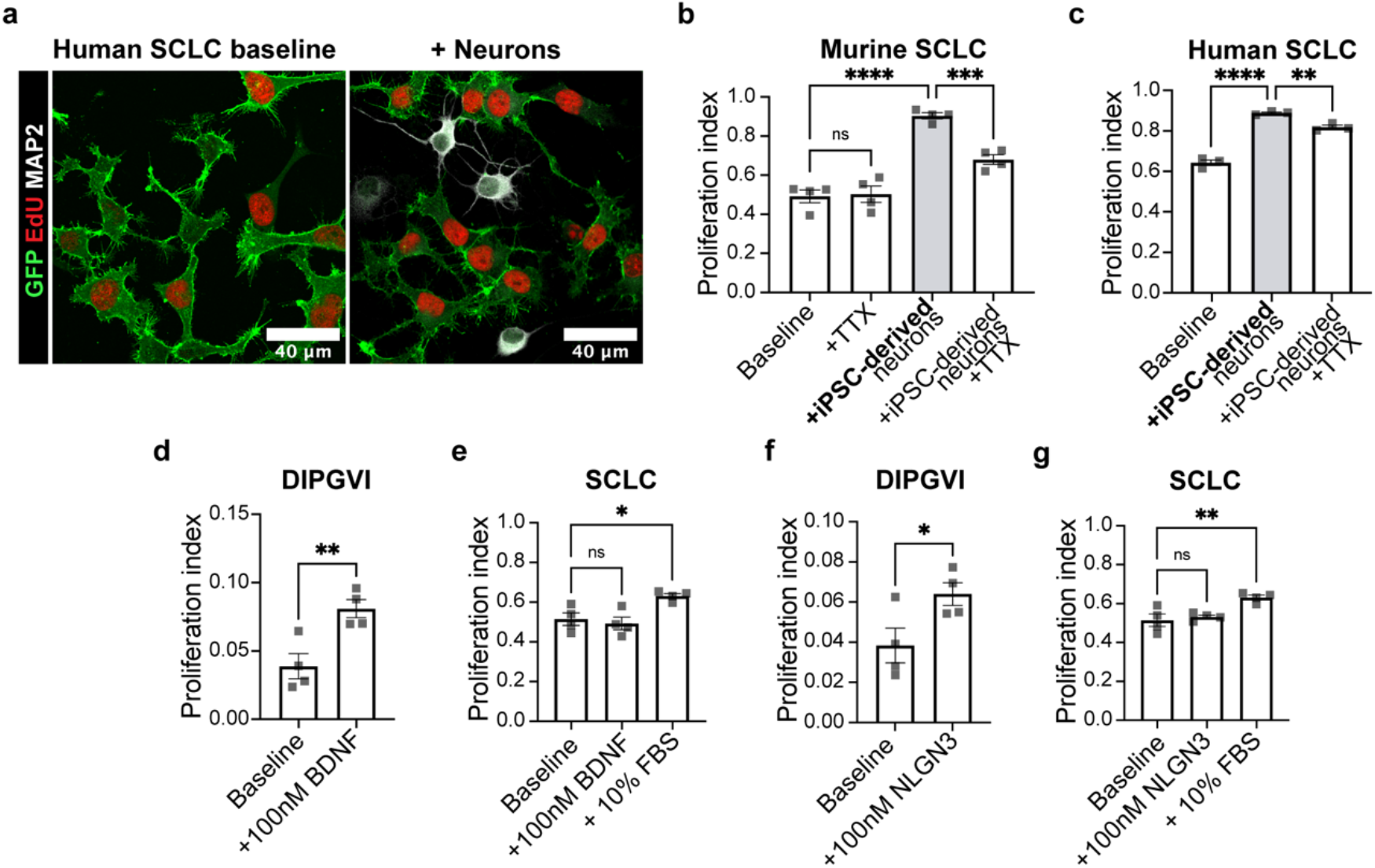
Neuron-SCLC co-cultures promote SCLC proliferation. **a**, Representative immunofluorescent images of human SCLC cells (H446; green) co-cultured with primary cortical neurons labeled with MAP2 (white). Proliferative cells are labeled with EdU (red). Scale bar = 40 μm. **b**, Quantification of proliferative index of murine SCLC cells (16T) co-cultured with human iPSC-derived neurons with or without addition of 1 μM TTX (n=4 coverslips per condition). **c**, as **b**, but for human SCLC (H446) cells (n=3 coverslips per condition). **d**, Quantification of proliferative index of patient derived glioma cells (SU-DIPGVI) at baseline or treated with 100 nM brain derived neurotrophic factor (BDNF; n=4 coverslips per condition). **e**, as in **d** but for SCLC cells. 10% FBS condition is used as positive control (n=4 coverslips per condition). **f**, Quantification of proliferative index of patient derived glioma cells (SU-DIPGVI) at baseline or treated with 100 nM neuroligin 3 (NLGN3; n=4 coverslips per condition). **g**, as in **f** but for SCLC cells. 10% FBS condition is used as positive control (n=4 coverslips per condition). Data are mean ± s.e.m.; analysis with 2-way ANOVA for **b, c**, two-tailed unpaired t-test for **d, e, f, g**. ****P<0.0001, ***P<0.001, **P< 0.01, *P< 0.05

**Extended Data Fig. 3.**
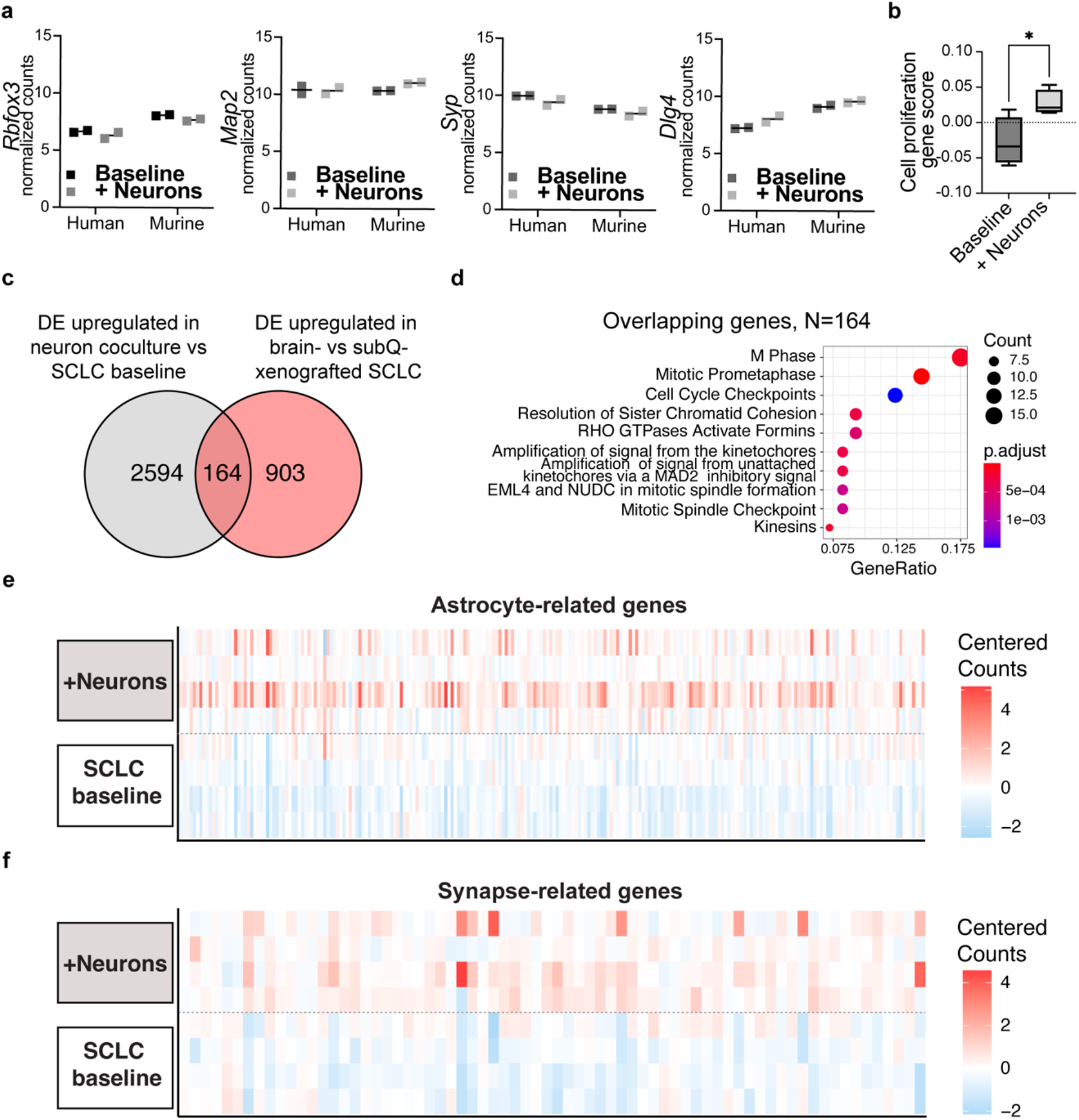
Transcriptional changes in SCLC after co-culture with neurons reflect increases in proliferation signature, astrocytic signature, and synaptogenic signature. **a**, Normalized counts of neuronal housekeeping genes in human and murine bulk RNAseq samples collected from SCLC cells at baseline (black) or isolated from neuronal co-culture (grey; n=4 independent samples per condition). **b**, Cell proliferation gene signature scores calculated from RNAseq of SCLC cells at baseline or isolated from neuronal co-cultures (n=4 independent samples per condition; for gene set used to generate gene signature see Extended Data Table 1). **c**, Venn diagram representing overlapping genes among (1) genes upregulated in SCLC cells co-cultured with neurons vs. in monoculture (grey) and (2) genes upregulated in SCLC cells allografted to the brain vs. subcutaneously (red). **d**, Gene Ontology (GO) terms enriched for in the 164 overlapping genes identified in **c. e**, Heatmap demonstrating centered counts of astrocyte-related genes from cultured murine and human SCLC cells at baseline or isolated from neuronal co-cultures (268 genes across n=4 samples from each condition; for the list of genes depicted see Extended Data Table 1). **f**, As in **e**, but for synapse-related genes (77 genes across n=4 samples from each condition; for the list of genes depicted see Extended Data Table 1). Data are plotted as median for **a**, box-and-whiskers plot for **b**; analyzed with two-way ANOVA for **a**, two-tailed unpaired t-test for **b**. *P< 0.05

**Extended figure 4.**
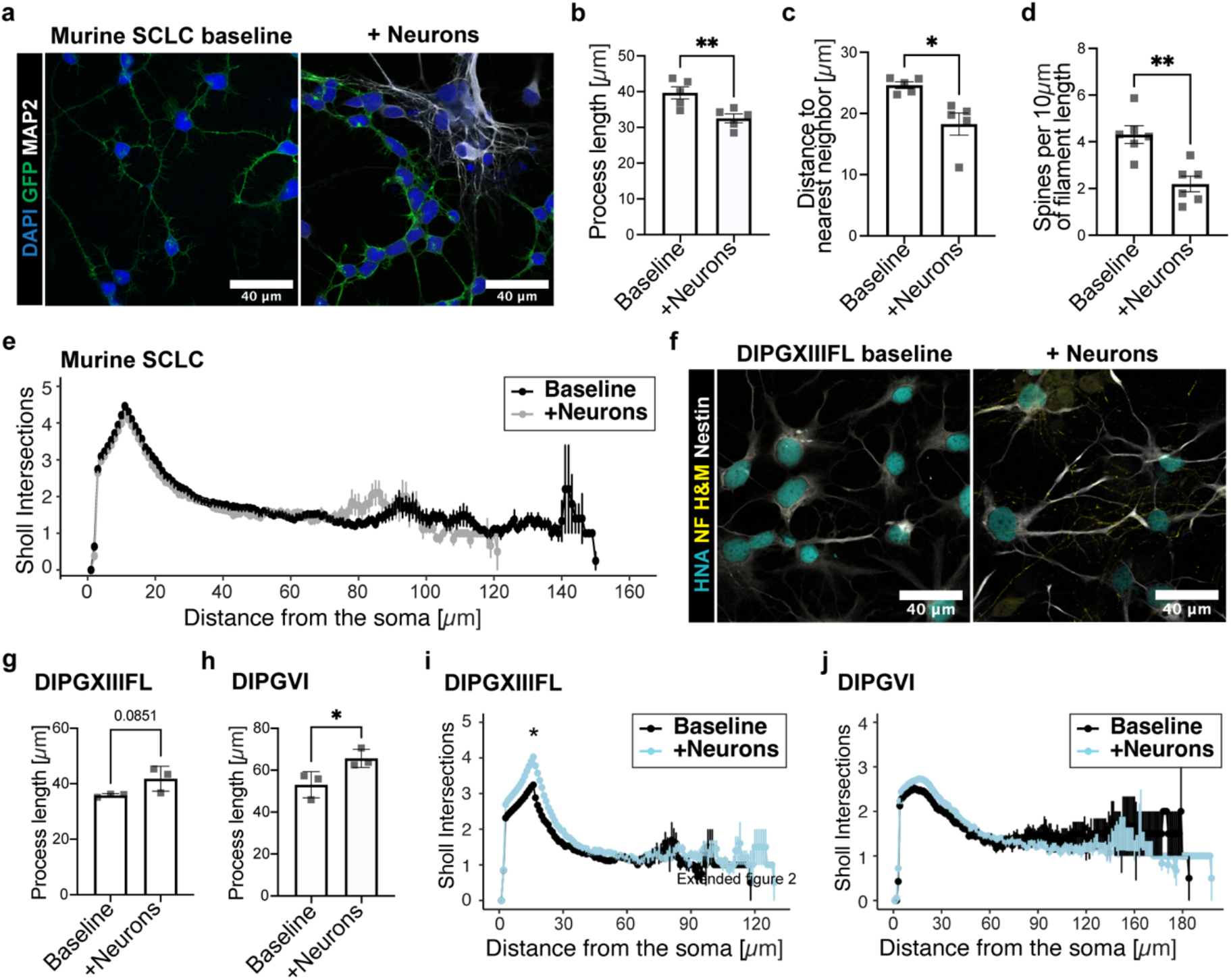
Neuronal co-culture induces a morphological shift in SCLC cells. **a**, Representative immunofluorescent images of human SCLC cells (16T-GFP, green) at baseline (left) or co-cultured (right) with primary neurons labeled with MAP2 (white). Scale bar = 40 μm. **b**, Quantification of median longest process length of SCLC cells at baseline or co-cultured with primary neurons (n=5 coverslips per condition). **c**, Quantification of median distance to nearest neighboring cell for SCLC cells at baseline or co-cultured with primary neurons. Data normalized to cell density (n=5 coverslips per condition). **d**, Quantification of the number of spine-like structures per 10 μm of SCLC cell process at baseline or co-cultured with primary neurons (n=6 coverslips per condition). **e**, Sholl plot depicting the mean number of intersections of tumor cell processes with concentric circles at increasing distances away from the tumor cell body for SCLC cells cultured alone (black) or in neuron co-cultures (grey) (n=1914 cells from 5 coverslips per condition). **f**, Representative immunofluorescent images of patient derived glioma cells (SU-DIPGXIII-FL; Nestin, white) at baseline (left) or co-cultured (right) with primary neurons labeled with neurofilament (yellow). Nuclei are labeled with human nuclear antigen (cyan). Scale bar = 40 μm. **g**, Quantification of median longest process length in glioma cells (SU-DIPGXIII-FL, a patient-derived culture of H3K27M-mutant diffuse midline glioma) at baseline or co-cultured with primary neurons (n=3 coverslips per condition). **h**, As in **g**, but for SU-DIPGVI glioma cells (n=3 coverslips per condition; a patient-derived culture of H3K27M-mutant diffuse midline glioma). **i**, As in **e**, but for SU-DIPGXIII-FL glioma cells cultured alone (black) or co-cultured with neurons in blue (n = 3356 cells from 3 coverslips per condition). **j**, As in **i**, but for SU-DIPGVI glioma cells (n=668 cells from 3 coverslips per condition). Data are mean ± s.e.m., two-tailed unpaired t-test for **b, c, d, g, h**. Sholl data analyzed using ANOVA with linear mixed-effects model fit using coverslip and cell id as random effects. **P< 0.01, *P< 0.05

**Extended figure 5.**
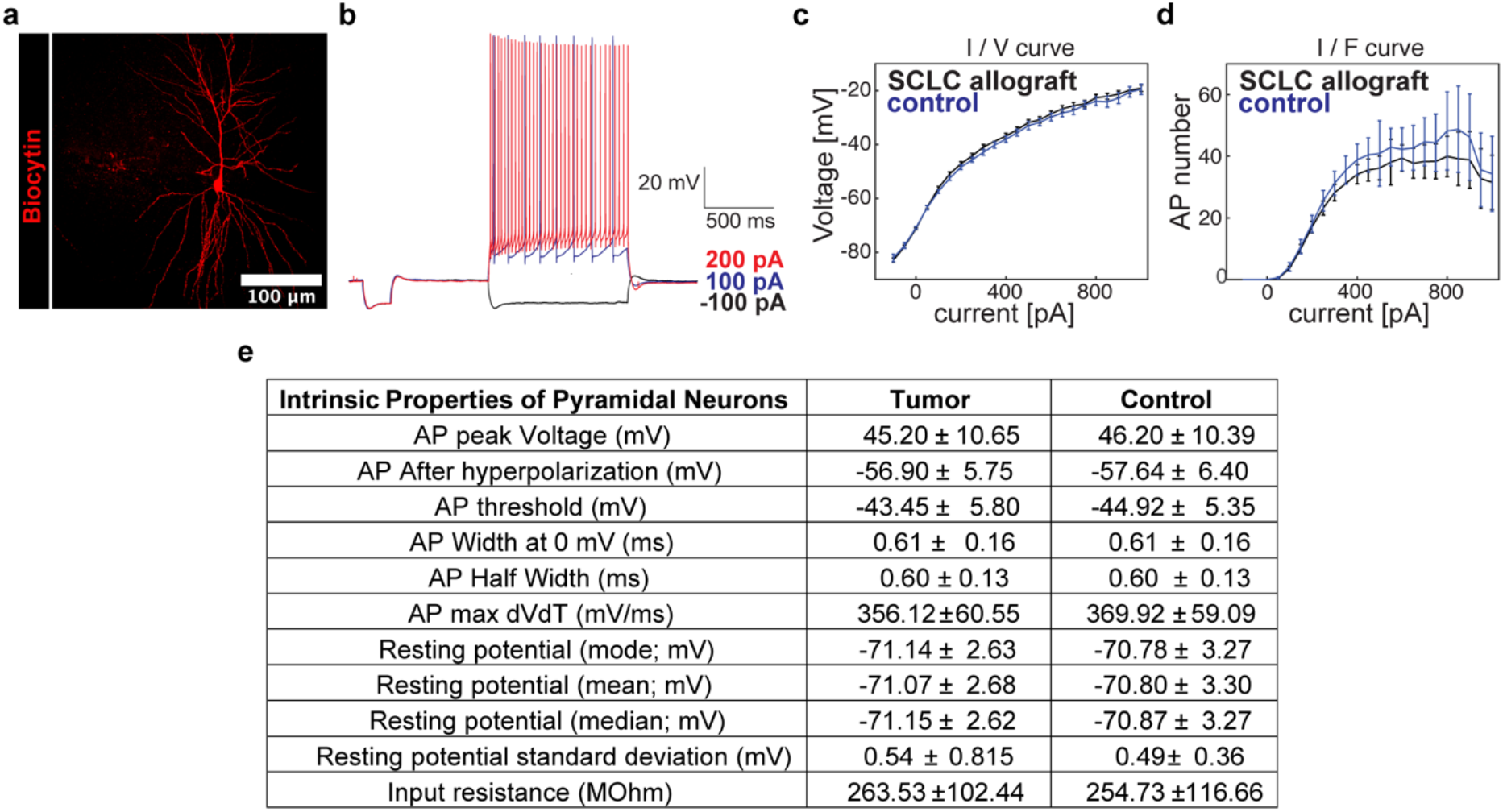
Cell intrinsic properties of neurons in and outside tumor regions. **a**, Biocytin (red)-filled pyramidal neuron in the area of the tumor cells. Scale bar = 100 μm. **b**, Representative current clamp currents and induced action potentials measured in pyramidal neurons in response to varying current injections (−100 pA, black; 100 pA, blue; 200 pA, red). **c**, Current to voltage relationship of action potentials in neurons from either the allograft (tumor-bearing) hippocampus or control contralateral hippocampus (n= 65 SCLC-associated, 32 control neurons). **d**, Current to action potential firing frequency relationship in neurons from either the allograft (tumor-bearing) hippocampus or control contralateral hippocampus (n= 65 SCLC-associated, 32 control neurons). **e**, Table listing cell intrinsic properties of pyramidal neurons from either the allograft (tumor-bearing) hippocampus or control contralateral hippocampus (n= 65 SCLC-associated, 32 control neurons). Data are mean ± s.e.m for **c, d, e**.

**Extended figure 6.**
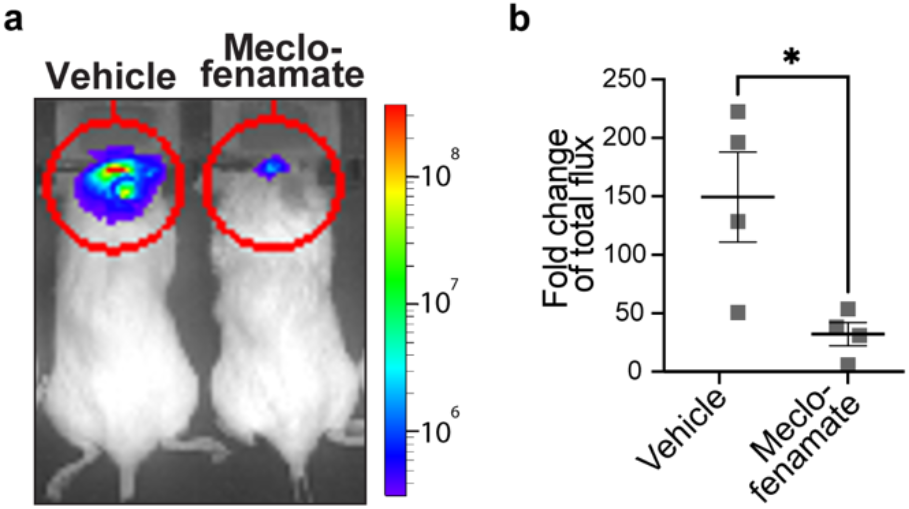
Blocking gap-junction mediated current inhibits growth of SCLC in the brain. **a**, Representative IVIS bioluminescence image of overall tumor burden of SCLC-luc allografts treated with vehicle or meclofenamate over a two-week period. Heat map, photon emission. **b**, Quantification of IVIS bioluminescence data from **a**, represented as fold change in total flux (n = 4 mice per group). Data are mean ± s.e.m., two-tailed unpaired t-test for **b**; *P< 0.05.

**Extended figure 7.**
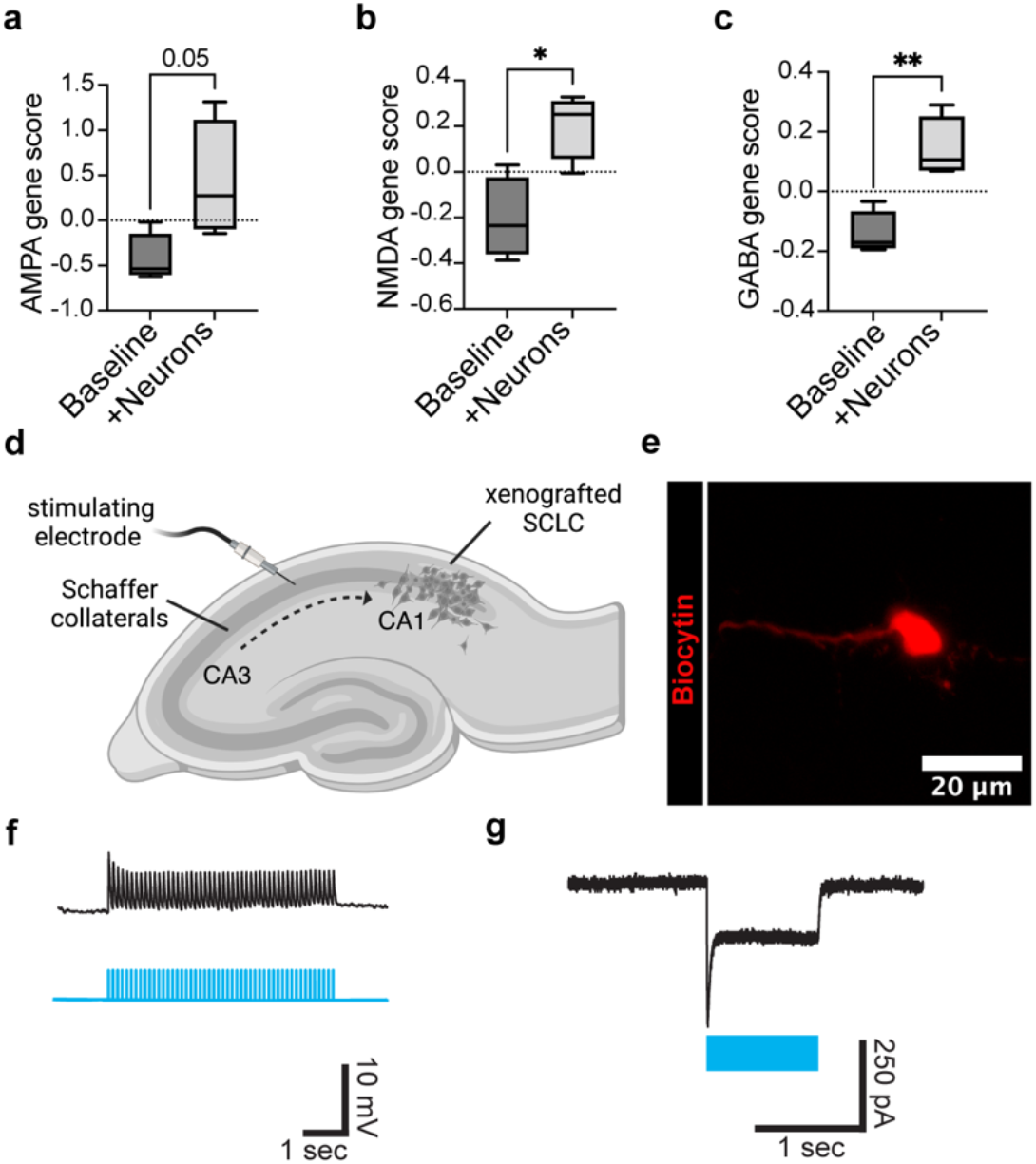
Gene signatures of SCLC cells in response to neuronal co-culture and electrical properties of SCLC cells. **a**, AMPA receptor gene signature scores calculated from RNAseq of SCLC cells (both human and murine) at baseline or isolated from neuronal co-cultures (n=4 samples per condition; for gene list used to generate transcriptomic signature see Extended Data Table 1). **b**, as in **a** for NMDA receptor gene signature score (n=4 samples per condition; see Extended Data Table 1). **c**, as in **a** for GABA receptor gene signature score (n=4 samples per condition; see Extended Data Table 1). **d**, Experimental paradigm for acute slice electrophysiology. GFP+ SCLC cells (grey) allografted in mouse hippocampus CA1 region with CA3 Schaffer collateral afferent stimulation. **e**, Biocytin-filled (red) SCLC cell allografted to CA1 region of mouse hippocampus. Scale bar = 20 μm. **f**, Whole cell patch-clamp voltage trace from ChR2-expressing SCLC cells (16T-ChR2) in response to blue light induced depolarization. Lower panel represents timing of blue light delivery. **g**, as in **f**, but demonstrating whole cell patch-clamp current trace from ChR2-expressing SCLC cells (16T-ChR2). Data are plotted as box-and-whisker plots, analysis with two-tailed unpaired t-test for **a-c**; **P< 0.01, *P< 0.05.

**Extended figure 8.**
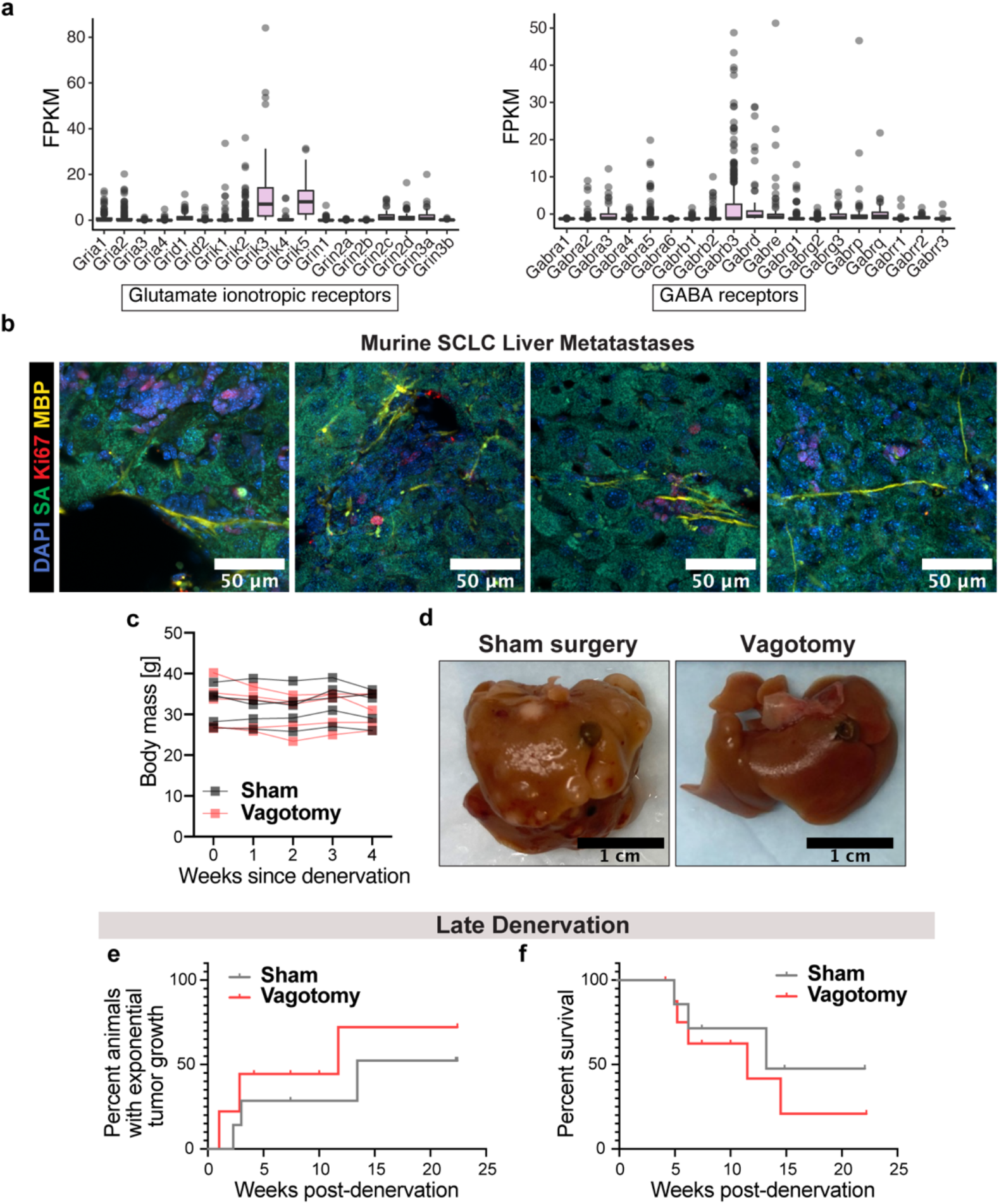
Denervation in genetic model of spontaneously forming SCLC. **a**, Quantification of glutamate and GABA receptor gene expression in human samples of primary SCLC (n=81, for gene list see Extended Data Table 1). **b**, Visualization of tumor innervation of liver metastasis in murine genetic model of SCLC. Streptavidin (SA, green) used to mark epithelial cells^67^; proliferative cells are labeled with Ki67 (red) to help identify tumors; MBP (yellow) is used to visualize nerve fibers. Scale bar = 50 μm. **c**, Body mass surveillance of animals up to 4 weeks following vagotomy or sham procedure (n=5 animals per group). **d**, Gross image of livers harvested at the endpoint of the experiment from the sham-operated (left) and denervated (right) animals. Scale bar = 1 cm. **e**, Time course of tumor growth as measured by IVIS bioluminescence imaging in sham-operated (grey) and denervated (vagotomy; red) animals that underwent surgery after initial tumor formation (n=7 sham and n=9 vagotomy animals from 2 independent cohorts). **f**, Kaplan–Meier survival curve for sham-operated (grey) and denervated (vagotomy; red) animals that underwent surgery after initial tumor formation (n=7 sham and n=9 vagotomy animals from 2 independent cohorts). Data analysis with Gehan-Breslow-Wilcoxon test for **e**, Log-rank (Mantel-Cox) test for **f**.

## Methods

### Mice and housing conditions

All *in vivo* experiments were conducted in accordance with protocols approved by the Brigham and Women’s Hospital Institutional Animal Care and Use Committee (IACUC) and Stanford University IACUC. Animals were housed according to the standard guidelines with free access to food and water in a 12 h light:12 h dark cycle.

For brain tumor allograft experiments, NSG mice (NOD-SCID-IL2R gamma chain-deficient, the Jackson Laboratory) were used. Male and female mice were used equally. According to the IACUC guidelines, signs of morbidity rather than maximal tumor volume was used as indication for termination of brain allograft animal experiments. Mice were euthanized if they exhibited signs of neurological disease or if they lost 15% or more of their body weight. For in-vivo optogenetic stimulation of the premotor circuit (M2), Thy1::ChR2;NSG or WT;NSG were used.

For lung tumor experiments, animals were euthanized when they exhibited signs of sickness behavior (dyspnea, abnormal gait or posturing, ill-groomed fur, etc.) or lost >15% of body weight. In these experiments, *Trp53*^*flox/flox*^, *Rb1*^*flox/flox*^, *p130*^*flox/flox*^, luciferase (luc) expressing genetic mouse models were used as described previously^62,68^. In these animals, lung tumors, and later distant metastases form spontaneously after intratracheal administration of Adeno-CMV-Cre (University of Iowa Vector Core, Iowa city, Iowa) at 2 months of age as described here^68^ and following a published protocol^69^.

### Intracranial allografts

All SCLC brain allografts were performed as previously described^4^. In brief, a single-cell suspension from cultured 16T-mGFP SCLC neurospheres was prepared in sterile HBSS immediately before surgery. Animals at postnatal day (P) 21–35 were anaesthetized with 1–4% isoflurane and placed in a stereotactic apparatus. The cranium was exposed via midline incision under aseptic conditions. 70,000 cells in 3 μL sterile HBSS were stereotactically injected into the M2 region of the cortex through a 31-gauge burr hole using a digital pump at infusion rate of 0.4 μL min^−1^ and 31-gauge Hamilton syringe. Stereotactic coordinates used were as follows: 0.5 mm lateral to midline, 1.0 mm anterior to bregma, −1.5 mm deep to cortical surface. At the completion of infusion, the syringe needle was allowed to remain in place for a minimum of 2 min, then withdrawn at a rate of 0.875 mm min^−1^ to minimize backflow of the injected cell suspension. For optogenetic experiments, tumors were allowed one week to engraft before implantation of the optogenetic ferrule. Allografts of 16T-mGFP-ChR2 SCLC cells were performed following the procedure as above with exception for the following alterations: 15,000 cells were injected in 1 μL sterile HBSS. Stereotactic coordinates used were as follows: 0.8 mm lateral to midline, 1.0 mm anterior to bregma, −1.2 mm deep to cortical surface.

### In-vivo optogenetic manipulation

For in-vivo optogenetic stimulation of M2 region of Thy1::ChR2;NSG or WT;NSG animals, a single stimulation paradigm was employed as previously described^1^. In brief, a fibre optic ferule was placed 1 week following and ipsilateral to the SCLC allografts. After 1 week to allow for recovery from the procedure, the animals were connected to a 100-mW 473-nm DPSS laser system with a mono fibre patch cord, which freely permits wakeful behavior of the mice. Pulses of light with approximately 4 mW measured output at tip of the patch cord were administered at a frequency of 20 Hz for periods of 30 s, followed by 90 s recovery periods, for a total session duration of 30 min. The animals were euthanized after 24 hours post stimulation, and brains were collected for histological analysis.

For in-vivo optogenetic depolarization of SCLC cells, ChR2–YFP (pLV-ef1-ChR2(H134R)-eYFP WPRE) construct (generated by Karl Deisseroth’s lab and placed in the piggyback transposon system by Minhui Su in the Monje lab) was lentivirally transduced into 16T SCLC cells, which were then allografted into premotor cortex (M2) following the procedure described above. A fibre optic ferule was implanted during the same surgery ipsilateral to the cell injection site at following coordinates: 0.8 mm lateral to midline, 1.0 mm anterior to bregma, −0.9 mm deep to cortical surface. At 1-week post-allograft and for 3 consecutive days, all animals were connected to the laser system to receive blue light or mock stimulations at a frequency of 10 Hz for periods of 30 s, followed by 90 s recovery periods, for a total session duration of 30 min. Mice were euthanized 24 h after the final (3rd) stimulation session.

### Immunohistochemistry

Immunohistochemistry of patient tissue samples was performed on formalin-fixed paraffin-embedded tissue sections per standard protocols including deparaffinization, antigen retrieval, incubation with primary antibody, and detection per the manufacturers’ instructions. The following antibodies were used: mouse anti-Ki-67 (Dako/Agilent), mouse anti-neurofilament (Ventana Roche; prediluted). Staining for Ki-67 was performed on a Leica Bond III automated stainer. Staining for neurofilament was performed on a Ventana Ultra automated stainer.

### Unilateral cervical vagotomy

Adult animals (4–5 months old) weighing over 17 grams were anaesthetized with 1-4% isoflurane through a nose cone in a supine position. Skin of the ventral surface of the neck was shaved and aseptically prepared according to IACUC guidelines. Under a dissection microscope, a 1 cm midline skin incision was made, and salivary glands revealed were separated with blunt dissection to expose the airways. The right vagal nerve was then carefully dissected from the carotid sheath and cut at the cervical level posterior to the pharyngeal branch. Sham animals underwent the same surgical procedure for blunt dissection of the vagus nerve but the latter was left intact. The animals were monitored till recovery from anesthesia for changes in heart rate and respiration and after that monitored bi-weekly for changes in weight, eating, drinking, and general activity.

### Bioluminescence imaging

For *in vivo* monitoring of tumor growth, bioluminescence imaging was performed using an IVIS imaging system (Xenogen). Animals were placed under 1-4% isofluorane anaesthesia and injected with luciferin substrate, then imaged in pronated position for intracranial allograft experiments or supine position for lung tumor animals. Baseline bioluminescence was used to randomize animals by a blinded investigator so that experimental groups contained an equivalent range of tumor sizes. For vagotomy experiments, each cohort of animals was imaged weekly for a total of 6 months.

### Survival studies

For survival studies, morbidity criteria used were either reduction of weight by 15% of initial weight, sickness behavior (dyspnea, abnormal gait or posturing, ill-groomed fur, etc.), or severe neurological motor deficits consistent with brainstem dysfunction (that is, hemiplegia or an incessant stereotyped circling behavior seen with ventral midbrain dysfunction). Kaplan–Meier survival analysis using log rank testing was performed to determine statistical significance.

### Histology

Animals with intracranial tumor allografts were anaesthetized with intraperitoneal avertin (tribromoethanol), then transcardially perfused with 20 ml of ice-cold PBS. Brains were fixed in 4% PFA overnight at 4°C, then cryoprotected in 30% sucrose, embedded in Tissue-Tek O.C.T. (Sakura) and sectioned in the coronal plane at 40μm using a sliding microtome. Lung tumor animals were perfused as above but also with 10 ml ice-cold 4% PFA. The lungs were inflated with 1-3 ml 2% UltraPure Low Melting Point Agarose (Invitrogen). Lungs and livers were fixed overnight at 4°C on a shaker, then transferred to 70% ethanol and sectioned at 15 μm for H&E and immunohistochemistry, or cryoprotected in 30% sucrose and sectioned at 150 μm on a cryotome for visualizing tumor innervation.

For immunofluorescence staining, the coronal sections were incubated in blocking solution (3% normal donkey serum, 0.3% Triton X-100 in TBS) at room temperature for 45 min, followed by an overnight incubation with primary antibodies in antibody diluent solution (1% normal donkey serum in 0.3% Triton X-100 in TBS) at 4 °C. On the next day, after a 5-min rinse in TBS, sections were incubated with DAPI (1μg/ml in TBS, Thermo Fisher) for 5 min, and rinsed again with TBS for 5 min. Afterwards, slices were incubated in secondary antibody solution at 4 °C overnight, then washed thrice in TBS and mounted with ProLong Gold Mounting medium (Life Technologies). For visualizing lung and liver tumor innervation with immunofluorescence, 150 μm thick sections were processed as above, except primary antibody incubation time was extended to 72 hours. Streptavidin Alexa Fluor 594 conjugate (Invitrogen) was used to visualized airway epithelium^67^. To quantify lung and liver tumor burden, 5 equidistant H&E sections from each organ were evaluated by a pathologist blinded to the experimental conditions to estimate percent of section area occupied by the tumor.

The following primary antibodies were used: chicken anti-GFP (Aves Labs, 1:500), rabbit anti-MAP2 (EMD Millipore, 1:500), mouse anti-NeuN (EMD Millipore, 1:500), rabbit anti-Ki67 (Abcam, 1:500), guinea pig anti-synapsin (Synaptic Systems, 1:500), rabbit anti-homer1 (Synaptic Systems, 1:500), mouse anti-neurofilament (Abcam, 1:500), mouse anti-nestin (Abcam, 1:1000), guinea pig anti-VChAT (Synaptic Systems, 1:200), mouse anti-TH (Abcam, 1:200), rat anti-MBP (Abcam, 1:200). The following secondary antibodies were used (all Jackson ImmunoResearch, 1:500): DyLight 405 Donkey Anti-Mouse IgG, Alexa Fluor 488 Donkey Anti-Chicken IgG, Alexa Fluor 488 Donkey Anti-Guinea Pig IgG, Alexa Fluor 488 Donkey Anti-Rabbit IgG, Alexa Fluor 594 Donkey Anti-Rabbit IgG, Alexa Fluor 647 Donkey Anti-Mouse IgG, Alexa Fluor 647 Donkey Anti-Rabbit IgG, Alexa Fluor 647 Donkey Anti-Rat IgG.

### Cell culture

The murine 16T SCLC line was derived from individual primary tumors from the lungs of Rb/p53 DKO mice. These cells are grown as neurospheres (unless otherwise stated) in 10% FBS medium consisting of DMEM (Invitrogen, Carlsbad, CA) and 1X liquid Antibiotic-Antimycotic (Invitrogen). The spheres were dissociated using TrypLE (Gibco) for seeding of *in vitro* experiments. Human H446 SCLC line was originally purchased from ATCC and cell identities were validated by Genetica DNA Laboratories using STR analysis. H446 cells are grown as attached cultures in RPMI (Thermo Fisher Scientific) with 10% FBS, 1x GlutaMax, and 1X liquid Antibiotic-Antimycotic (Invitrogen). All cultures are monitored by short tandem repeat (STR) fingerprinting for authenticity throughout the culture period and mycoplasma testing was routinely performed.

### Co-culture of SCLC cells with primary murine neurons

Neurons were isolated from the brains of CD1 animals using the ‘Neural Tissue Dissociation Kit - Postnatal Neurons’ (Miltenyi), followed by the ‘Neuron Isolation Kit, Mouse’ (Miltenyi) per manufacturer’s instructions. After isolation, 300,000 neurons were plated onto circular glass coverslips (Electron Microscopy Services) pre-treated for 20min at 37°C with poly-L-lysine (Sigma) and then 3 hours at 37°C with 5 μg/mL mouse laminin (Thermo Fisher). Neurons are cultured in BrainPhys neuronal medium (Stemcell Technologies) supplemented with 1x glutamax (Invitrogen), pen/strep (Invitrogen), B27 supplement (Invitrogen), BDNF (10ng/mL; Shenandoah), and GDNF (5ng/mL; Shenandoah), TRO19622 (5μM; Tocris), beta-mercaptoethanol (1X, Gibco), and 2% fetal bovine serum. Half of the medium was replenished on DIV 1 and UFDU was added at 1μM. This was repeated at DIV 3. On DIV 5, half of the medium was replaced with serum-free medium in the morning. In the afternoon, the medium was again replaced with half serum-free medium containing 75,000 SCLC cells dissociated from neurospheres or attached cultures with TrypLE. Tumor cells were cultured with neurons for 24 hours and then fixed with 4% PFA for 20 minutes at room temperature and stained for immunofluorescence analysis as described below.

### Co-culture of SCLC cells with human iPSC-derived neurons

Induced pluripotent stem cell lines were obtained from the Brigham and Women’s Hospital NeuroHub Core Facility and all permissions were received for use (from BWH NeuroHub Core and Rush Alzheimer’s Disease Center’s Biospecimen Distribution Committee). Induced neurons were generated from BR33 iPSCs as previously described^70^. Briefly, iPSCs were plated in mTeSR1 media at a density of 95K cells/cm2 on Matrigel-coated plates for viral transduction. Viral plasmids were obtained from Addgene (plasmids #19780, 52047, 30130). FUdeltaGW-rtTA was a gift from Konrad Hochedlinger (Addgene plasmid # 19780). Tet-O-FUW-EGFP was a gift from Marius Wernig (Addgene plasmid # 30130). pTet-O-Ngn2-puro was a gift from Marius Wernig (Addgene plasmid # 52047). Lentiviruses were obtained from Alstem with ultrahigh titers (∼3 × 10^9^) and used at the following concentrations: pTet-ONGN2-puro: 0.1 μl/ 50K cells; Tet-O-FUW-eGFP: 0.05 μl/ 50K cells; Fudelta GW-rtTA: 0.11ul/50K cells. Transduced cells were dissociated with Accutase and plated onto Matrigel-coated plates at 50,000 cells/cm2 in StemFlex media (day 0). On day 1, media was changed to KSR media with doxycycline (2 ug/ml, Sigma). Doxycyline was maintained in the media for the remainder of the differentiation. On day 2, media was changed to 1:1 KSR: N2B media with puromycin (5 ug/ml, GIBCO). Puromycin was maintained in the media throughout the differentiation. On day 3, media was changed to N2B media + 1:100 B27 supplement (Life Technologies), and puromycin (10 ug/ml). From day 4 on, cells were cultured in NBM media + 1:50 B27 + BDNF, GDNF, CNTF (10 ng/ml, Peprotech). Around D12-D-15, SCLC cells were added a ratio of 1:3 (cancer:neuron). Cultures were monitored over the next 10 days in the case of MEA recordings. For histological analyses, cultures were fixed and analyzed 24 hours after co-culture in the case of proliferation assays, and 5 days after co-culture in the case of synapse quantification.

### Induced neuron protocol media

#### KSR media

Knockout DMEM, 15% KOSR, 1x MEM-NEAA, 55 uM beta-mercaptoethanol, 1x GlutaMAX (Life Technologies).

#### N2B media

DMEM/F12, 1x GlutaMAX (Life Technologies), 1x N2 supplement B (StemCell Technologies, Inc.), 0.3% dextrose (D-(+)-glucose, Sigma).

#### NBM media

Neurobasal medium, 0.5x MEM-NEAA, 1x GlutaMAX (Life Technologies), 0.3% dextrose (D-(+)-glucose, Sigma).

### RNA sequencing from co-culture experiments

Fastq files were aligned to the human (hg19) and mouse (grcm38) genome using hisat2 (v2.1.0). Feature counting was performed using rsem (v1.3.0). Following alignment, all analyses were performed in R (v4.1.0). Centered count values plotted in heatmaps were calculated by first log transforming TPM count matrices followed by subtracting mean gene expression values from each sample’s normalized gene expression value.

Differential expression comparing cells with and without neuron co-culture was performed on raw count matrices using DESeq2 (v1.32.0; fitType = parametric, size factor estimation = ratio). To control for batch effects, replicate ID was included in the design matrix for DESeq2 (∼Replicate + Condition). The Wald test was used for significance testing. Genes with a logFC greater than 1 and an adjusted p-value (FDR) less than 0.05 were considered significantly differentially expressed. To test for enriched gene ontology terms, the enrichGO function from the R package clusterProfiler (v4.0.5) was used to identify over-represented terms (pAdjustMethod=“BH”, qvalueCutoff=0.05, minGSSize = 10, maxGSSize = 500).

Gene signature scores were computed as described previously^71^. Given a gene set *G* for an NMF program or reference gene set, a score *S* was generated to quantify the relative expression of *G* within each sample *i*. This was calculated as the mean centered expression *CE* of genes in *G* compared to the centered expression of a background gene set *bG*, determined as the 100 genes with most similar aggregate expression level to each gene in *G*. The final score for a given sample *i* and gene set *G* is calculated as 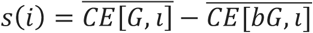.

### Synaptic puncta staining and visualization

For immunohistochemistry, fixed coverslips were incubated in blocking solution (3% normal donkey serum, 0.3% Triton X-100 in TBS) at room temperature for 1 h. Primary antibodies guinea pig anti-synapsin1/2 (1:500; Synaptic Systems), rabbit anti-Homer1 (1:500; Synaptic Systems), or mouse anti-neurofilament (1:500; Abcam) in 0.3% Triton X-100 in TBS and incubated overnight at 4 °C. Samples were then rinsed three times in TBS and incubated in secondary antibody solution (Alexa 488 donkey anti-guinea pig IgG; Alexa 594 donkey anti-rabbit IgG, and Alexa 647 donkey anti-mouse IgG, all at 1:500 (Jackson Immuno Research)) in antibody diluent solution at 4 °C overnight. Coverslips were rinsed three times in TBS and mounted with ProLong Gold Mounting medium (Life Technologies). Images were collected using a 63X oil-immersion objective on a Zeiss LSM800 confocal microscope and processed with Airyscan. Colocalization of puncta was quantified as previously described^4^.

### Multielectrode array recordings

All multielectrode array (MEA) recordings were taken and analyzed using the Axion Biosystems platform. Prior to culturing, 6-well Axion plates were coated with Poly-L-Lysine and laminin. D4 iNs (created as previously described^70^) were then thawed and plated in neurobasal media, at 100,000 cells per well. For the initial week prior to co-culture, iNs were subjected to a half media change every 3-4 days. Doxycycline and puromycin treatment were stopped after D10 to allow for the addition of tumor cells. On D14-D15, SCLC cells were added at 30,000 cells/well MEA plates. MEA plates were recorded every day for 10 minutes for up to 2 weeks days after co-culture. All spike numbers and amplitude were assessed using proprietary Axion Software.

### EdU incorporation assay

EdU staining was performed on glass coverslips in 24-well plates which were precoated with poly-l-lysine (Sigma) and 5 μg/mL mouse laminin (Thermo Fisher). Neurosphere cultures were dissociated with TrypLE and plated onto coated slides with 10 μM of EdU. After 24 hours the cells were fixed with 4% paraformaldehyde in PBS for 20 minutes and then stained using the Click-iT 594 EdU kit and protocol (Invitrogen) with or without additional antibody staining and mounted using Prolong Gold mounting medium (Life Technologies). Proliferation index was determined by quantifying the fraction of EdU labeled cells/GFP labeled cells using confocal microscopy at 40x magnification.

### Cell morphology analysis

Confocal images of SCLC cells cultured in the presence or absence of neurons were analyzed using the autopath mode of filament tracer function in Imaris 9.9.0. The generated 3D renderings of cell processes were described by quantifying the length of the longest process, the distance of the cell body to the nearest neighbor, and the number of spine-like structures per unit of length of the process.

### Mouse drug treatment studies

For drug studies, NSG mice were allografted as above with 16T-GFP-luc cells and randomized to treatment group by a blinded investigator. One week post-allograft, tumor-bearing mice were treated with systemic administration of meclofenamate sodium (20 mg kg−1; Selleck Chemicals; formulated in 10% DMSO in PBS) via intraperitoneal injection for two weeks (5 days per week). Controls were treated with an identical volume of vehicle. Bioluminescence imaging was performed before treatment and every 4 days thereafter using an IVIS imaging system (Xenogen) under isoflurane anaesthesia.

### Electrophysiology

Brain slices were obtained from 2- to 4-month-old mice (both male and female) using standard techniques. In brief, mice were anesthetized by isoflurane inhalation and perfused transcardially with ice-cold artificial cerebrospinal fluid (ACSF) containing (in mM) 125 NaCl, 2.5 KCl, 25 NaHCO_3_, 2 CaCl_2_, 1 MgCl_2_, 1.25 NaH_2_PO_4_ and 25 glucose (295 mOsm/kg). Brains were blocked and transferred into a slicing chamber containing ice-cold ACSF. Coronal slices of hippocampus were cut at 300 μm thickness with a Leica VT1000s vibratome in ice-cold ACSF, transferred for 10 min to a holding chamber containing choline-based solution consisting of (in mM): 110 choline chloride, 25 NaHCO_3_, 2.5 KCl, 7 MgCl_2_, 0.5 CaCl_2_, 1.25 NaH_2_PO_4_, 25 glucose, 11.6 ascorbic acid, and 3.1 pyruvic acid at 34°C, and then transferred to a secondary holding chamber containing ACSF at 34°C for 10 min and subsequently maintained at room temperature (20–22°C) until use. For electrophysiology recordings, individual brain slices were transferred into a recording chamber, mounted on an upright microscope (Olympus BX51WI) and continuously superfused (2–3 ml min^−1^) with ACSF warmed to 32–34°C by passing it through a feedback-controlled in-line heater (SH-27B; Warner Instruments). Cells were visualized through a 60X water-immersion objective with either infrared differential interference contrast optics or epifluorescence to identify GFP+ SCLC cells. For whole cell voltage clamp recording, patch pipettes (2–4 MΩ) pulled from borosilicate glass (G150F-3, Warner Instruments) were filled with internal solution containing (in mM) 135 CsMeSO_3_, 10 HEPES, 1 EGTA, 3.3 QX-314 (Cl− salt), 4 Mg-ATP, 0.3 Na-GTP, 8 Na_2_-phosphocreatine (pH 7.3 adjusted with CsOH; 295 mOsm·kg−1). Spontaneous currents were recorded for 5 minutes at a holding potential of – 70 mV. To record evoked currents of GFP+ SCLC cells, the membrane voltages were clamped at –70 mV or 0 mV, extracellular stimulation was performed with a stimulus isolation unit (MicroProbes, ISO-Flex) with bipolar electrodes (100 μm apart, PlasticOne) placed 100-400 μm perpendicularly away from recording cells (0.1ms, 100-200uA, delivered at 10 second intervals). Pyramidal cells in allografted and contralateral sites of hippocampus were recorded in whole cell current clamp to study intrinsic properties. Patch pipettes were filled with internal solution containing (in mM) 135 KMeSO3, 10 HEPES, 1 EGTA, 4 Mg-ATP, 0.4 Na-GTP, 8 Na_2_-phosphocreatine (pH 7.3 adjusted with KOH; 295 mOsm·kg−1). At −70mV, currents were injected for 1000ms at 50pA steps from −100 to 1000pA. Intrinsic properties of hippocampus pyramidal cells were analyzed with custom program written for Matlab. All data were acquired with a MultiClamp 700B amplifier (Molecular Devices) and digitized at 10 kHz with a National Instruments data acquisition device (NI USB-6343).

### Calcium imaging

For calcium imaging, the genetically encoded calcium indicator GCaMP6s was lentivirally transduced into murine SCLC 16T (pLV-ef1-GCaMP6s-P2A-nls-tdTomato). In this case, SCLC cells containing the GCaMP6s reporter can be identified using the tdTomato nuclear tag. These cells were isolated and grafted into the CA1 region of the hippocampus as described above. Two-photon calcium imaging experiments were performed using Prairie Ultima XY upright two-photon microscope for tissue slices equipped with an Olympus LUM Plan FI W/IR-2 40× water immersion objective. The temperature of the perfusion media, ACSF as described above, was kept at 30°C, and perfused through the system at rate of 2 ml min−1. Excitation light was provided at a wavelength of 920 nm through a tunable Ti:Sapphire laser (Spectra Physics Mai Tai DeepSee) to allow for excitation of both tdTomato and GCaMP6s. The actual laser power reaching the scanhead for each scope is dynamically controlled by Pockels cells via software interface. Pockels cell were set at 10 for all experiments, and PMTs were set at 800 for each channel. For these settings, power at back aperture of the objective was approximately 30 mW at 920 nm. The wavelength ranges for the emission filters were PMT1: 607 nm centre wavelength with 45 nm bandpass (full-width at half-maximum) and PMT2: 525-nm centre wavelength with 70-nm bandpass (full-width at half-maximum). Recordings were made at 0.65 frames per second (∼1.5 Hz) for about 30 min in the case of spontaneous activity and 10 min in the case of response to periodic electrical stimulation. Cells were identified via the expression for the nuclear tdTomato tag and were only imaged in the area of interest, specifically in the CA1 region of the hippocampus. Similar to the electrophysiology paradigm, for neuronal stimulation experiments, the electrode was placed in the hippocampus to stimulate the neuronal inputs originating in CA3. For electrical stimulation, approximately 20 μA over 200 μs was delivered to local axons using a stimulating bipolar microelectrode. For all inhibitor experiments, inhibitors were directly diluted into the ACSF perfusion media at desired concentration, oxygenated, and delivered to the slices through the perfusion system.

### Calcium imaging analysis

Quantitative fluorescence intensity analysis was done on calcium transients that were reliably evoked by axonal stimulation. To determine the effect of tetrodotoxin on the calcium transients in response to electrical stimulation of the CA3 Schaffer collaterals, the field of cells were stimulated three times in 1-min intervals to ensure synaptic connectivity. TTX (500 μM) was then perfused into the slices and the stim was repeated on the same field of cells to gauge direct effect of TTX on stimulation response. For analysis, ROIs of each responding nucleus were set and ΔF_max_/F_0_ (maximum difference in fluorescence intensity normalized to background fluorescence) measurements were determined before and after TTX treatment.

### Statistical analyses

Statistical tests were conducted using Prism v9.1.0 (GraphPad) software unless otherwise indicated. Gaussian distribution was confirmed by the Shapiro-Wilk normality test. For parametric data, unpaired two-tailed Student’s t-test or one-way ANOVA with Tukey’s post hoc tests to examine pairwise differences were used as indicated. Paired two-tailed Student’s t-tests were used in the case of same cell or same animal experiments (as in electrophysiological recordings and vagotomy experiments). For non-parametric data, a two-sided unpaired Mann-Whitney test was used as indicated, or a one-tailed Wilcoxon matched pairs signed rank test was used for same-cell experiments. Two-tailed log rank analyses were used to analyze statistical significance of Kaplan-Meier survival curves. In all box-and-whiskers plots, whiskers indicate minimum and maximum value, box extends to 25^th^ and 75^th^ percentile, the line in the middle is plotted at the median. In all violin plots, lines are drawn at the median and quartiles.

## Data availability

RNA sequencing of SCLC cells isolated from neuronal co-cultures will be made available on Gene Expression Omnibus (GEO) prior to publication. All other data are available in the manuscript or from the corresponding author upon reasonable request. Source data will be uploaded with the final version of the manuscript.

## Code availability

Sources for all code used have been provided, no custom code was created for this manuscript.

## Author conflicts of interest

M.M. holds equity in MapLight Therapeutics and Syncopation Life Sciences. M.M. was previously in the SAB for Cygnal Therapeutics.

## Acknowledgements

Research reported in this publication was supported by the NIH (grants CA252001 to H.S.V.; R01NS092597, DP1NS111132; P50CA165962, R01CA258384, U19CA264504 to M.M.; and CA231997 to J.S.), Glaucoma Research Foundation (to H.S.V), Charles Hood Foundation (to H.S.V), ChadTough Defeat DIPG (to M.M. and H.S.V.), Virginia and D.K. Ludwig Fund for Cancer Research (to M.M.), Cancer Research UK (to M.M.), Robert J. Kleberg, Jr. and Helen C. Kleberg Foundation (to M.M.), McKenna Claire Foundation (to M.M.), Damon Runyon Cancer Research Foundation fellowship (F.Q.), Stanford Medical Scholars Research Program (S.S.). We kindly thank Dr. Kerriann Casey for reviewing murine lung and liver histology.

## Author contributions

S.S., M.M. and H.S.V. designed experiments. S.S. and H.S.V. conducted and analyzed experiments. S.S. contributed to all *in vitro* and *in vivo* data collection and analysis. W.W. and B.L.S. contributed to electrophysiology experiments. K.G., E.C., and E.C.P. contributed to co-culture and MEA experiments and analysis. A.M.T performed histological staining of human tissue. B.Y. and P.J.W. contributed to *in vivo* optogenetic surgeries. Y.L. and M.A.K. contributed to the vagotomy surgeries and strategy. F.Q. and J.S. provided SCLC mice and RNAseq from brain and subcutaneously allografted cells. J.L. and M.G.F. contributed to bioinformatic analyses. All authors contributed to manuscript editing. M.M. and H.S.V. wrote the manuscript and supervised all aspects of the work. H.S.V. conceived of the project.

## References

1. Venkatesh, H. S. et al. Neuronal Activity Promotes Glioma Growth through Neuroligin-3 Secretion. Cell 161, 803–16 (2015).

2. Venkatesh, H. & Monje, M. Neuronal Activity in Ontogeny and Oncology. Trends in cancer 3, 89–112 (2017).

3. Pan, Y. et al. NF1 mutation drives neuronal activity-dependent initiation of optic glioma. Nature 594, 277–282 (2021).

4. Venkatesh, H. S. et al. Electrical and synaptic integration of glioma into neural circuits. Nature 573, 539–545 (2019).

5. Taylor, K. R. et al. Glioma synapses recruit mechanisms of adaptive plasticity. bioRxiv 2021.11.04.467325 (2021). doi:10.1101/2021.11.04.467325

6. Venkataramani, V. et al. Glutamatergic synaptic input to glioma cells drives brain tumour progression. Nature 573, 532–538 (2019).

7. Zeng, Q. et al. Synaptic proximity enables NMDAR signalling to promote brain metastasis. Nature 573, 526–531 (2019).

8. Magnon, C. et al. Autonomic nerve development contributes to prostate cancer progression. Science 341, 1236361 (2013).

9. Renz, B. W. et al. Cholinergic Signaling via Muscarinic Receptors Directly and Indirectly Suppresses Pancreatic Tumorigenesis and Cancer Stemness. Cancer Discov. 8, 1458– 1473 (2018).

10. Zhao, C.-M. et al. Denervation suppresses gastric tumorigenesis. Sci. Transl. Med. 6, 250ra115 (2014).

11. Hayakawa, Y. et al. Nerve Growth Factor Promotes Gastric Tumorigenesis through Aberrant Cholinergic Signaling. Cancer Cell 31, 21–34 (2017).

12. Renz, B. W. et al. β2 Adrenergic-Neurotrophin Feedforward Loop Promotes Pancreatic Cancer. Cancer Cell 33, 75–90.e7 (2018).

13. Peterson, S. C. et al. Basal cell carcinoma preferentially arises from stem cells within hair follicle and mechanosensory niches. Cell Stem Cell 16, 400–12 (2015).

14. Kamiya, A. et al. Genetic manipulation of autonomic nerve fiber innervation and activity and its effect on breast cancer progression. Nat. Neurosci. 22, 1289–1305 (2019).

15. Saloman, J. L. et al. Ablation of sensory neurons in a genetic model of pancreatic ductal adenocarcinoma slows initiation and progression of cancer. Proc. Natl. Acad. Sci. U. S. A. 113, 3078–83 (2016).

16. Sloan, E. K. et al. The sympathetic nervous system induces a metastatic switch in primary breast cancer. Cancer Res. 70, 7042–52 (2010).

17. Stopczynski, R. E. et al. Neuroplastic changes occur early in the development of pancreatic ductal adenocarcinoma. Cancer Res. 74, 1718–27 (2014).

18. Sabari, J. K., Lok, B. H., Laird, J. H., Poirier, J. T. & Rudin, C. M. Unravelling the biology of SCLC: implications for therapy. Nat. Rev. Clin. Oncol. 14, 549–561 (2017).

19. Rudin, C. M., Brambilla, E., Faivre-Finn, C. & Sage, J. Small-cell lung cancer. Nat. Rev. Dis. Prim. 7, 3 (2021).

20. Wang, Y. et al. Exploration of spatial distribution of brain metastasis from small cell lung cancer and identification of metastatic risk level of brain regions: a multicenter, retrospective study. Cancer Imaging 21, 41 (2021).

21. Riihimäki, M. et al. Metastatic sites and survival in lung cancer. Lung Cancer 86, 78–84 (2014).

22. Park, K.-S. et al. Characterization of the cell of origin for small cell lung cancer. Cell Cycle 10, 2806–2815 (2011).

23. Semenova, E. A., Nagel, R. & Berns, A. Origins, genetic landscape, and emerging therapies of small cell lung cancer. Genes Dev. 29, 1447–62 (2015).

24. Pan, J., Yeger, H. & Cutz, E. Innervation of Pulmonary Neuroendocrine Cells and Neuroepithelial Bodies in Developing Rabbit Lung. Journal of Histochemistry & Cytochemistry 52, (2004).

25. Lauweryns, J. M. & Van Lommel, A. Ultrastructure of nerve endings and synaptic junctions in rabbit intrapulmonary neuroepithelial bodies: a single and serial section analysis. J. Anat. 151, 65–83 (1987).

26. Lauweryns, J. M. & Peuskens, J. C. Neuro-epithelial bodies (neuroreceptor or secretory organs?) in human infant bronchial and bronchiolar epithelium. Anat. Rec. 172, 471–81 (1972).

27. Anderson, N. E., Rosenblum, M. K., Graus, F., Wiley, R. G. & Posner, J. B. Autoantibodies in paraneoplastic syndromes associated with small-cell lung cancer. Neurology 38, 1391– 1391 (1988).

28. Yang, D. et al. Axon-like protrusions promote small cell lung cancer migration and metastasis. bioRxiv 726026 (2019). doi:10.1101/726026

29. Onganer, P. U., Seckl, M. J. & Djamgoz, M. B. A. Neuronal characteristics of small-cell lung cancer. Br. J. Cancer 93, 1197–1201 (2005).

30. Dong, A., Zhang, J., Chen, X., Ren, X. & Zhang, X. Diagnostic value of ProGRP for small cell lung cancer in different stages. J. Thorac. Dis. 11, 1182–1189 (2019).

31. Oc, B. et al. Serum neuron-specific enolase is a useful tumor marker for small cell lung cancer. Cancer 65, (1990).

32. Akoun, G. M., Scarna, H. M., Milleron, B. J., Bénichou, M. P. & Herman, D. P. Serum neuron-specific enolase. A marker for disease extent and response to therapy for small-cell lung cancer. Chest 87, 39–43 (1985).

33. Qu, F. et al. Neuronal mimicry generates an ecosystem critical for brain metastatic growth of SCLC. bioRxiv 2021.08.10.455426 (2021). doi:10.1101/2021.08.10.455426

34. Ko, J., Winslow, M. M. & Sage, J. Mechanisms of small cell lung cancer metastasis. EMBO Mol. Med. 13, e13122 (2021).

35. NCI-H446 [H446] - HTB-171 | ATCC. Available at: https://www.atcc.org/products/htb-171. (Accessed: 28th November 2022)

36. Venkatesh, H. S. & Monje, M. Neuronal Activity in Ontogeny and Oncology. Trends in Cancer (2017). doi:10.1016/j.trecan.2016.12.008

37. Cahoy, J. D. et al. A transcriptome database for astrocytes, neurons, and oligodendrocytes: a new resource for understanding brain development and function. J. Neurosci. 28, 264–78 (2008).

38. Doyle, J. P. et al. Application of a translational profiling approach for the comparative analysis of CNS cell types. Cell 135, 749–62 (2008).

39. Lovatt, D. et al. The transcriptome and metabolic gene signature of protoplasmic astrocytes in the adult murine cortex. J. Neurosci. 27, 12255–66 (2007).

40. Lim, J. S. et al. Intratumoural heterogeneity generated by Notch signalling promotes small-cell lung cancer. Nature 545, 360–364 (2017).

41. Denny, S. K. et al. Nfib Promotes Metastasis through a Widespread Increase in Chromatin Accessibility. Cell 166, 328–342 (2016).

42. Lee, J.-H. et al. Astrocytes phagocytose adult hippocampal synapses for circuit homeostasis. Nature 590, 612–617 (2021).

43. Takano, T. et al. Chemico-genetic discovery of astrocytic control of inhibition in vivo. Nature 588, 296–302 (2020).

44. Ws, C., Nj, A. & c, E. Astrocytes Control Synapse Formation, Function, and Elimination. Cold Spring Harb. Perspect. Biol. 7, (2015).

45. Araque, A., Parpura, V., Sanzgiri, R. P. & Haydon, P. G. Tripartite synapses: glia, the unacknowledged partner. Trends Neurosci. 22, 208–15 (1999).

46. Chung, W.-S. et al. Astrocytes mediate synapse elimination through MEGF10 and MERTK pathways. Nature 504, 394–400 (2013).

47. Sibille, J., Pannasch, U. & Rouach, N. Astroglial potassium clearance contributes to short-term plasticity of synaptically evoked currents at the tripartite synapse. J. Physiol. 592, 87–102 (2014).

48. Wolpert, F. et al. Postoperative progression of brain metastasis is associated with seizures. Epilepsia 63, e138–e143 (2022).

49. Azary, S. et al. Incidence of Seizure and Associated Risk Factors in Patients in the Medical Intensive Care Unit (ICU) at Memorial Sloan Kettering Cancer Center (MSK) from 2016-2017. J. Intensive Care Med. 37, 1312–1317 (2022).

50. Avila, E. K. & Graber, J. Seizures and epilepsy in cancer patients. Curr. Neurol. Neurosci. Rep. 10, 60–7 (2010).

51. Lee, M. H., Kong, D.-S., Seol, H. J., Nam, D.-H. & Lee, J.-I. Risk of seizure and its clinical implication in the patients with cerebral metastasis from lung cancer. Acta Neurochir. (Wien). 155, 1833–7 (2013).

52. Buckingham, S. C. et al. Glutamate release by primary brain tumors induces epileptic activity. Nat. Med. 17, 1269–74 (2011).

53. John Lin, C. C. et al. Identification of diverse astrocyte populations and their malignant analogs. Nat. Neurosci. (2017). doi:10.1038/nn.4493

54. Yu, K. et al. PIK3CA variants selectively initiate brain hyperactivity during gliomagenesis. Nature 578, 166–171 (2020).

55. Krishna, S. et al. Glioblastoma remodeling of neural circuits in the human brain decreases survival. bioRxiv (2021).

56. Chen, Q. et al. Carcinoma–astrocyte gap junctions promote brain metastasis by cGAMP transfer. Nature 533, 493–498 (2016).

57. Osswald, M. et al. Brain tumour cells interconnect to a functional and resistant network. Nature 528, 93–8 (2015).

58. Venkataramani, V. et al. Glioblastoma hijacks neuronal mechanisms for brain invasion. Cell 185, 2899–2917.e31 (2022).

59. Yabumoto, Y. et al. Expression of GABAergic system in pulmonary neuroendocrine cells and airway epithelial cells in GAD67-GFP knock-in mice. Med. Mol. Morphol. 41, 20–7 (2008).

60. Barrios, J. et al. Pulmonary Neuroendocrine Cells Secrete γ-Aminobutyric Acid to Induce Goblet Cell Hyperplasia in Primate Models. Am. J. Respir. Cell Mol. Biol. 60, 687–694 (2019).

61. George, J. et al. Comprehensive genomic profiles of small cell lung cancer. Nature 524, 47–53 (2015).

62. Schaffer, B. E. et al. Loss of p130 accelerates tumor development in a mouse model for human small-cell lung carcinoma. Cancer Res. 70, 3877–83 (2010).

63. Gazdar, A. F. et al. The Comparative Pathology of Genetically Engineered Mouse Models for Neuroendocrine Carcinomas of the Lung. J. Thorac. Oncol. 10, 553–564 (2015).

64. Kwon, M. & Berns, A. Mouse models for lung cancer. Mol. Oncol. 7, 165–177 (2013).

65. Rudin, C. M. et al. Molecular subtypes of small cell lung cancer: a synthesis of human and mouse model data. Nat. Rev. Cancer 19, 289–297 (2019).

66. De Virgiliis, F. & Di Giovanni, S. Lung innervation in the eye of a cytokine storm: neuroimmune interactions and COVID-19. Nat. Rev. Neurol. 16, 645–652 (2020).

67. Scott, G. D., Blum, E. D., Fryer, A. D. & Jacoby, D. B. Tissue optical clearing, three-dimensional imaging, and computer morphometry in whole mouse lungs and human airways. Am. J. Respir. Cell Mol. Biol. 51, 43–55 (2014).

68. Denny, S. K. et al. Nfib Promotes Metastasis through a Widespread Increase in Chromatin Accessibility. Cell 166, 328–342 (2016).

69. DuPage, M., Dooley, A. L. & Jacks, T. Conditional mouse lung cancer models using adenoviral or lentiviral delivery of Cre recombinase. Nat. Protoc. 4, 1064–72 (2009).

70. Lagomarsino, V. N. et al. Stem cell-derived neurons reflect features of protein networks, neuropathology, and cognitive outcome of their aged human donors. Neuron 109, 3402–3420.e9 (2021).

71. Gojo, J. et al. Single-Cell RNA-Seq Reveals Cellular Hierarchies and Impaired Developmental Trajectories in Pediatric Ependymoma. Cancer Cell 38, 44–59.e9 (2020).

